# CharacTERT: A Machine Learning Tool for Classifying hTERT Missense Variants

**DOI:** 10.64898/2026.05.18.725793

**Authors:** Georgina Becerra Parra, Qisheng Pan, Yoochan Myung, Stephanie Portelli, Niles E. Nelson, Joanne L. Dickinson, Sionne E.M. Lucas, Jessica K. Holien, Tracy M. Bryan, David B. Ascher

## Abstract

Missense mutations in *TERT*, the gene encoding the human telomerase catalytic subunit hTERT, are associated with Telomere Biology Disorders (TBDs). Experimentally elucidating the effects of all possible missense variants would be time-consuming and technically challenging. Moreover, current computational predictors are not hTERT-specific and primarily rely on sequence information, failing to capture the complex biological and structural context of the telomerase enzyme. In this work, we developed three machine learning models integrating both sequence- and structure-based features to account for the biological mechanisms of hTERT. Compared to state-of-the-art methods, our best-performing models achieved a higher Matthew’s Correlation Coefficient of 0.88 on ClinVar and gnomAD curated variants and demonstrated robust sensitivity (0.75) on a dataset curated according to guidelines from the American College of Medical Genetics and Genomics and Association for Molecular Pathology (ACMG/AMP). Feature interpretation highlighted hTERT residue conservation and changes in hydrophobic and weak polar interactions as critical determinants of pathogenicity. Finally, *in silico* saturation mutagenesis was performed to present a mutational landscape of *TERT*, available in a user-friendly web server, CharacTERT, which could offer valuable insights into the molecular mechanisms driving TBDs, aid in early diagnosis, as well as guide personalized treatment strategies. CharacTERT is freely available at https://biosig.lab.uq.edu.au/charactert/.

## Introduction

Telomere biology disorders (TBDs) encompass a broad spectrum of rare genetic conditions that affect telomere maintenance mechanisms (1–3). Since their discovery, these disorders have been linked to missense mutations in at least seventeen different genes, including *TERT*, the gene encoding the catalytic subunit of the enzyme telomerase, which is crucial for preserving telomere length in most organisms (4).

Telomerase is a ribonucleoprotein that plays a key role in maintaining telomere length by reverse transcription of an integral RNA template, named hTR in humans (5). Telomeres are nucleoprotein complexes located at the end of linear chromosomes. They primarily protect chromosomal ends from deoxyribonucleases (DNases) and prevent their fusion, thereby maintaining genome stability (6). In most normal human somatic cells, telomerase activity is minimal or absent, leading to gradual telomere shortening. This natural process is associated with aging and tissue degeneration and serves as a tumour-suppressive mechanism by limiting cellular proliferation (7). Consequently, *TERT* expression is tightly regulated and generally restricted to prenatal development, germ cells, and stem cells (8,9). However, germline mutations in *TERT*, which are frequently missense mutations, can result in accelerated telomere shortening, contributing to the development of TBDs (10). *TERT* mutations can display highly variable penetrance within families; TBDs also demonstrate marked anticipation, i.e. an increase in severity of the phenotype and earlier onset of disease in subsequent generations, due to inheritance of shorter telomeres with each generation. These factors often make it challenging to assign pathogenicity to newly identified variants in *TERT* and other TBD genes, which complicates management decisions and can delay treatment (11).

In humans, the canonical hTERT protein sequence consists of 1132 residues, organized into four main functional domains, each playing a critical role in the process of telomere extension: the telomerase essential N-terminal (TEN) domain, the telomerase RNA-binding domain (RBD), the reverse transcriptase (RT) domain, and the C-terminal extension (CTE) domain (see Figure 1A and Figure 1B). These protein domains, along with the 451-nucleotide RNA component hTR (shown in Figure 1C and Figure 1D), encoded by the gene *TERC,* coordinate together to sustain telomere length and genome stability (12–15).

**Figure 1:**
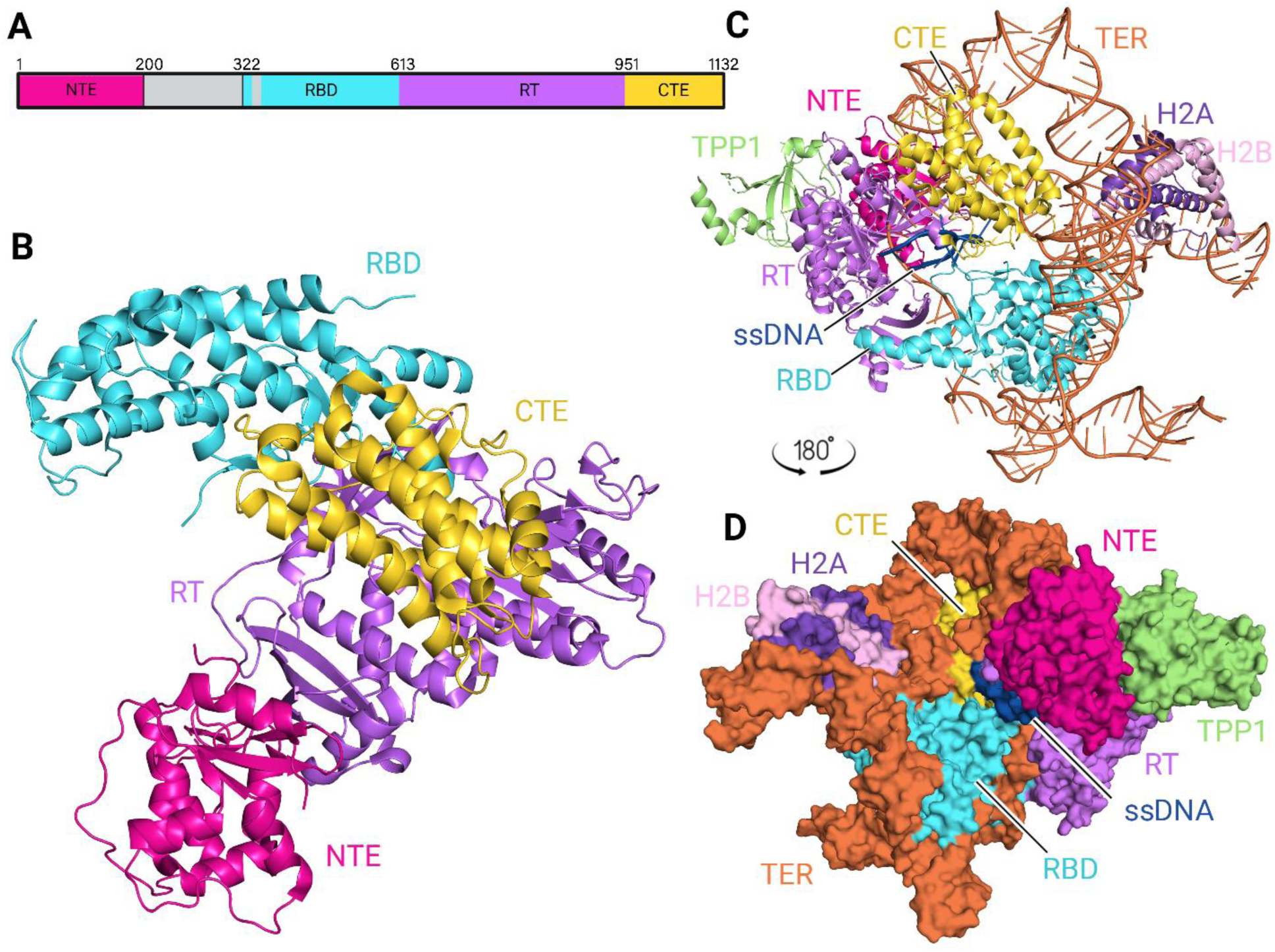
Structure of the telomerase catalytic subunit hTERT with telomeric DNA, RNA, and associated proteins. Domain organization of hTERT in the protein sequence (**A**) and in the PDB structure 7QXA (13) (**B**). Each domain is represented by different colours, named TEN or telomerase essential N-terminal domain (pink), RBD or RNA-binding domain (cyan), RT or reverse transcriptase domain (purple), and CTE or C-terminal extension domain (yellow). The complete PDB structure 7QXA includes telomerase RNA labelled hTR (orange) and TERT binding partners, named shelterin protein TPP1 (green), histone H2A (dark purple), histone H2B (light pink), and ssDNA or single stranded DNA (navy) (**C-D**).

The TEN domain contributes to TERT’s ability to add multiple telomere repeats by binding to TPP1 (encoded by the gene *ACD*), a shelterin component that stabilizes the enzyme’s position on the telomere by interacting with a specific RT domain region named TRAP (16,17). The RBD domain directly binds to hTR, the telomerase RNA component that provides a template for telomere synthesis. The RT domain serves as the catalytic core, driving the addition of nucleotide repeats to the telomere end, while the CTE domain is essential for anchoring the RNA-DNA complex, thereby enabling telomerase to synthesize multiple consecutive TTAGGG repeats without dissociating, a process known as repeat addition processivity (18,19). Together, these domains support the sequential steps required for hTERT’s function in addition of multiple TTAGGG repeats to telomere ends.

Disease-associated germline missense variants in *TERT* are found across these structural domains, and disruptions within each domain can lead to impaired telomerase activity (20), contributing to the spectrum of TBD phenotypes, including dyskeratosis congenita, idiopathic pulmonary fibrosis, liver disease, bone marrow failure and haematological malignancy. Recent advancements in sequencing technologies (21–23) have led to the identification of a wealth of *TERT* genetic variants. However, the evaluation of the pathogenic effects of hTERT missense variants remains challenging. Current experimental validation techniques and *in silico* tools have limitations. While experimental approaches are time-consuming, expensive, and technically challenging with limited throughput (11), available computational predictors are not hTERT*-*specific, making it difficult to achieve high predictive performance and account for the complex biological context of the protein (24). These limitations are reflected in the ClinVar database (25,26), where 85% of hTERT variants are considered variants of uncertain significance (VUS) (see Figure S1). Despite TERT’s critical role in cancer, cellular aging, and disease, no computational methods specifically designed to evaluate the impact of hTERT variants have been previously reported.

In this work, we harnessed both sequence- and structure-based features to identify pathogenic drivers of human hTERT, following a similar pipeline to previous works (27–29). Our models were developed using reported variants available in ClinVar (25,26), gnomAD (30), and the Telomerase database (20) and tested using a clinical dataset curated using the American College of Medical Genetics and Genomics/Association for Molecular Pathology (ACMG/AMP) criteria for pathogenicity (31,32). Statistical comparisons showed that disease-causing variants were more likely to be associated with changes in the number of hydrophobic or weak polar interactions, consequently disrupting hTERT function. Moreover, comparative analyses against state-of-the-art methods demonstrated that our models significantly outperformed these tools, particularly in predicting ACMG/AMP-curated variants across several metrics, suggesting potential clinical utility. In our work, we propose potential pathogenic molecular drivers of hTERT, which can guide earlier genetic disease diagnosis and the development of novel gene therapy approaches. Additionally, we developed a user-friendly web server, CharacTERT, that provides a prediction for the pathogenicity of all possible missense variants in hTERT, providing an invaluable resource to advance the understanding and diagnosis of *TERT*-associated disease.

## Method

The aim of this project was to develop an hTERT-specific pathogenicity predictor by identifying the features of the protein that are most likely to predict whether a variant is deleterious, enhancing the understanding of hTERT association with disease and providing a tool for clinical genetic laboratories to use in classifying newly identified VUS in *TERT*. To achieve this, we followed a structure-based variant analysis pipeline reported in our previous work (27–29), whose workflow is shown in Figure 2.

**Figure 2:**
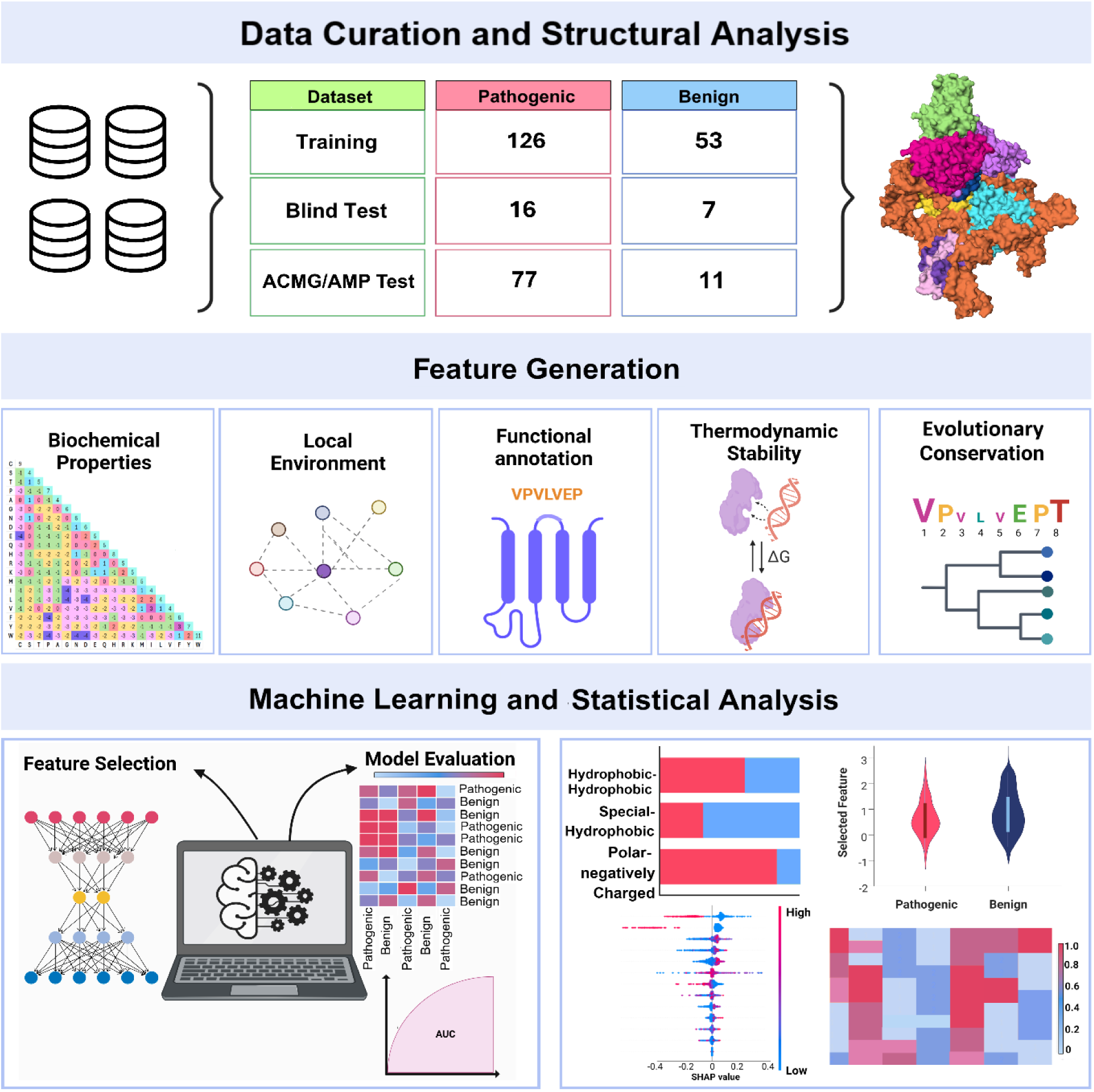
Pipeline to predict pathogenic missense variants of hTERT. Annotated missense variants were collected from multiple databases and subsequently distributed into two different sets: training and blind test. Structural information was obtained from the PDB structure 7QXA (13). After data and structure curation, the variants were characterized using several features, including biochemical properties, local environment, functional annotation, thermodynamic stability and conservation scores. Finally, statistical comparisons and machine learning analysis were implemented to assess potential risk factors leading to pathogenic variants in *TERT*.

## Data curation

### (a) Training and blind test data

We collated TERT missense variants from the databases ClinVar (25,26), gnomAD v4.1.0 (30) (n=124), and the Arizona State University Telomerase Database (20) (n=62) in February 2024. By integrating variants from these databases, we aimed to collect a comprehensive dataset, referred to herein as the TERT manually curated dataset, that captured the diversity of TERT variants, such as their clinical, population, and molecular relevance.

For ClinVar data, germline missense variants with at least one curation as “Pathogenic” or “Likely pathogenic” were considered pathogenic in this dataset, while those with a ClinVar curation of “Benign” or “Likely benign” were considered benign. Variants with conflicting interpretations between these two classifications were excluded from the datasets. It is worth noting that some variants received their classification based solely on a single curation from Invitae as “Likely Pathogenic” or “Likely Benign”, with no other non-VUS curations. An additional 62 variants were sourced from the Arizona State University Telomerase Database; since all these variants have been reported in the literature in patients with TBDs, they were considered pathogenic for this study. After removing redundant entries and variants with conflicting labels across the different databases, our final dataset consisted of 59 benign and 127 pathogenic variants (Tables S6-S7).

To ensure consistency with our structural analysis, variants located in the linker region between the TEN and RBD domains were excluded from this dataset, as this region was absent in the experimental structure used for feature generation.

### (b) Independent clinical validation set based on ACMG/AMP guidelines

An independent dataset of TERT missense variants meeting ACMG/AMP criteria for pathogenicity was curated by us for a separate study assessing the performance of current multi-gene trained in silico predictors of pathogenicity (Nelson et al., unpublished data), using a previously published methodology (11) for application of ACMG/AMP criteria. During this process, the PP3/BP4 criteria (i.e. available computational evidence) was excluded from variant classification since that was the parameter under study. To enhance the generalization of our models, redundant entries were removed by comparing this dataset with our training and blind test sets. This step ensured that the models were not biased by repeated variants and improved their ability to be generalized to unseen data. Moreover, 17 variants occurring at the same residue positions as those in our primary dataset were removed to minimize performance bias due to data leakage of position-specific features. This strategy ensured that the ACMG/AMP-based dataset included only unique variants, each with distinct residue positions and local environments. Notably, variants located in the linker region between the TEN and RBD domains were included in this dataset. Therefore, the final ACMG/AMP-based dataset consisted of 11 benign and 77 pathogenic variants (Table S8).

### (c) Experimental literature variants

In addition to the previously described datasets, we manually curated 30 TERT missense variants with published experimental functional data in February 2025, to further assess the predictive performance, generalizability, and robustness of the saturation mutagenesis predictions of our best-performing model (Table S9). These variants were identified through clinical screening, but at the time there was insufficient clinical or population evidence to classify them based on the ACMG/AMP guidelines. To maintain consistency with the ACMG/AMP-based dataset and minimize performance bias due to data leakage from position-specific features, variants occurring at the same positions as those in the TERT manually curated dataset were excluded from this set.

Functional classification of these variants as defective or normal was based on a systematic evaluation of experimental data, using predefined criteria reflecting telomerase activity, processivity, and/or cellular function. Variants were classified as functionally defective if they exhibited impaired telomerase function, meeting at least one of the following criteria: (i) telomerase activity or processivity <75% in a “direct” (i.e. non-PCR) activity assay (DTA), prioritizing cell-based telomerase reconstitution over in vitro reconstitution in rabbit reticulocyte lysates (RRL) in cases of discrepancy; (ii) activity <50% in a PCR-based Telomere Repeat Amplification Protocol (TRAP) assay, as long as conflicting results from DTA or telomere elongation assay (TEA) were not present; (iii) TEA or immortalization efficiency below 75%; or (iv) efficiency of telomerase recruitment to telomeres <75%. The DTA was prioritized over the TRAP assay due to the demonstrated low specificity of the latter assay (11).

Conversely, variants were classified as functionally normal using criteria that depend on their domain location. For those in the TEN (1–193) or IFD-TRAP (721–815) domains (33,34), functional normality required telomerase activity and processivity measured in a DTA ≥75%, prioritizing cell-based assays over RRL reconstitution where discrepancies existed, in combination with either telomere recruitment efficiency ≥75% or TEA or immortalization efficiency ≥75%. Variants located outside these domains were classified as functionally normal if either telomerase activity (DTA) and processivity were ≥75%, prioritizing cell-based assays over RRL in cases of discrepancy, or TEA or immortalization efficiency was ≥75%.

## Curation of available structures

Eight experimental full-length hTERT structures were available on the RCSB Protein Data Bank (PDB) as of February 2024 (35). We selected a protein complex (PDB ID: 7QXA) experimentally determined via cryo-electron microscopy to a resolution of 3.20 Å, consisting of hTERT, the telomerase RNA subunit, a telomeric DNA substrate oligonucleotide, and the interacting proteins histone H2A and H2B and telomere-binding protein TPP1 (encoded by gene *ACD*), to capture the biological context of hTERT’s functional environment (13) (see Figure 1C and Figure 1D).

## Biophysical and biochemical annotation of variants

We computationally characterized the effects of hTERT missense variants on its structure and function using five properties: biochemical properties and sequence-based information, local environment, domain annotation, predicted protein thermodynamic stability and interactions, and conservation.

### (a) Biochemical properties and sequence-based information

Single amino acid changes can significantly impact the biochemical properties of a protein region, potentially disrupting its function (36,37). To assess these impacts on hTERT function, multiple sequence alignment (MSA)-based biochemical properties from AAindex2 and 3 databases (38) were extracted including all the conservation-based substitution matrices PAM and BLOSUM (39–42). While these matrices are rooted in conservation-based substitution preferences, they also provide critical information on the potential biochemical effects of amino acid substitutions, such as hydrophobicity, charge complementarity, and hydrogen bonding tendencies.

### (b) Local environment

The residue environment of a variant site is essential to understand the structural and functional consequences of each variant. To achieve a multi-dimensional characterization of the hTERT variant environment, structural descriptors were generated from three complementary features:

#### (i) Local structural properties

Temperature factor, which measures the dynamic motion or positional uncertainty of an atom within a crystal structure, relative solvent accessibility, and residue depth of the variant site were computed using Biopython. Secondary structure types and torsion angles psi and phi were computed using the Dictionary of Secondary Structure of Proteins (DSSP) program (43). In addition, the disordered regions of hTERT were annotated using IUPRED (v.2 and v.3) and ANCHOR tools (44).

#### (ii) Residue interactions

We first used MODELLER (45,46) to model the mutated hTERT structures for each variant in the TERT manually curated dataset. Arpeggio (47) was then used to capture wild-type residue interactions, as well as their changes upon variant within a radius of 6 Å, including hydrophobic, weak polar, and proximal interaction counts.

#### (iii) Graph-based signatures

Graph-based signatures were computed using mCSM-suite (48). The signatures capture atomic distance patterns at the variant site by modelling the atom of different pharmacophores as nodes and their contacts with other atoms within a certain distance cutoff as edges. By setting a cutoff step of 0.5 Å and a distance interval between 1 and 12 Å, we systematically characterised the variant environment of hTERT. Pharmacophore changes of eight different types of atoms were computed, which were appended to the signatures to provide additional physicochemical context. Additionally, signatures of atom pairs were generated without distinguishing between different atom classes, ensuring the structural and chemical context of the variant site was captured in terms of functional group behaviours, rather than specific atom identities.

### (c) Domain and PTM site annotation

Domain information of human hTERT was extracted from UniProt (49). Variants were annotated based on their location within the functional domains (TEN, RBD, RT, and CTE) in a one-hot encoded manner, a binary representation method where 1 indicated the presence of the variant in that domain and 0 elsewhere. ActiveDriverDB (50) was used to identify whether the variant site could have potential post-translational modifications, such as phosphorylation, methylation, or glycosylation.

### (d) Predicted impact of variants on protein stability and interactions

Variants in hTERT can affect protein stability, compromising its essential role in telomere maintenance and potentially leading to TBDs. To characterise these effects, DDMut (51) and DynaMut (52,53) were used to compute the predicted change of protein stability upon variant. Additionally, as hTERT carries out its function by interacting with its telomeric DNA substrate, the telomerase RNA subunit, and other binding proteins such as TPP1, destabilizing variants may impair these associations, further hindering telomere elongation. To assess these, mCSM-NA (54) was used to study the predicted effect of variant on the interaction between hTERT and DNA and RNA chains, while mCSM-PPI and mCSM-PPI2 (48,55) were used to assess potential changes in protein-protein interactions (see Figure 1C-1D).

### (e) Conservation scores

Highly evolutionarily conserved regions tend to show lower tolerance to mutations, often resulting in significant disease implications. Therefore, evolutionary conservation scores for each residue were calculated using ConSurf (56), GEMME (57) and Envision (58) tools. To assess population conservation, missense tolerance ratios (MTRs) were obtained from MTR-viewer (v.1 and v.2) (59,60) and MTR-3D (61) resources using different sliding windows (21, 31 and 41 codons), a spatial radius of 5Å, and the PDB structure 7QXA. These MTRs can help identify specific regions of the protein sequence that exhibit low tolerance to missense mutations, which tend to be crucial for maintaining protein function and structural integrity.

## Statistical analysis

Before implementing machine learning, we conducted a statistical analysis to identify potential pathogenic drivers of hTERT. Given the limited dataset size, the two-tailed Wilcoxon rank-sum test (62,63) was implemented for statistical comparison of the different features that may not follow a normal distribution.

We initially focused on analysing features that were expected to exhibit significant differences (p-value < 0.05) between the two classes: pathogenic and benign. These features included changes in protein stability, protein-nucleic acid interactions, and protein-protein contacts, all of which are directly related to the biological function of hTERT. Following this, we calculated p-values for 295 features and selected the most significant ones for further discussion.

## Supervised machine learning

### (a) Model development

The original dataset was randomly divided into training (Pathogenic: 114; Benign: 53) and blind test (Pathogenic: 13; Benign: 6) sets, in a 90:10 split (Tables S6-S7). The training set was further divided into a sub-training set (90% of the training set, approximately 80% of the original dataset; Pathogenic: 100; Benign: 47), and a validation set (10% of the training set, approximately 10% of the original dataset; Pathogenic: 14; Benign: 6) for greedy feature selection. To reduce redundancy and enhance model generalization, variants occurring at the same residue position were grouped together in either training, blind test, or validation sets. Moreover, the distribution of pathogenic and benign labels was balanced across the different sets.

To assess how the position and domain of hTERT missense variants influenced pathogenicity prediction, three different machine learning models were developed. To ensure robust model performance and prevent overfitting, k-fold cross-validation with k-values of 10, 4, and 3 was employed to train each model, respectively (64). Variants in each fold were grouped based on either the position they were occurring in or the domain in which they were located, depending on the model being developed. During this process, we ensured that each fold carefully maintained consistent proportions of both pathogenic and benign variants by using StratifiedGroupFold in sci-kit learn.

To further assess the generalization ability of our models, an independent ACMG/AMP-based dataset was used for evaluation. Predictive performance was measured using different metrics, including Matthew’s correlation coefficient (MCC), accuracy, sensitivity, and specificity.

In this work, we adopted the machine learning approach detailed in previous studies (26–28), testing eleven algorithms on the training set using default parameters: AdaBoost (n_estimators=300), Decision Tree (random_state=1), ExtraTrees (n_estimators=300), Gradient Boosting (n_estimators=300), Extreme Gradient Boosting (n_estimators=300), K-nearest neighbour (k=3), Logistic Regression (random_state=1), Multilayer Perceptron (random_state=1), Gaussian Naive Bayes (random_state=1), Random Forest (n_estimators=300), and Support Vector Machines (kernel =’rbf’), using the scikit-learn package (v1.0.2) (65).

### (b) Feature selection and interpretation

Among the eleven machine learning algorithms, the three models with the best predictive performance were further optimized using greedy feature selection, based on the MCC metric. This method employs a forward sequential feature selection approach that iteratively identifies the best new feature to enhance model performance, which in this study was assessed using the MCC metric. MCC provides a balanced measure of model performance, particularly in datasets with class imbalances, making it a more reliable indicator of predictive capacity in our work. The process starts with an empty set of features, and at each step, the feature that maximizes the cross-validated score is added. This continues until a predetermined number of features is reached (66). Greedy feature selection was performed using the sub-training set, while the validation set was used to assess the performance of the selected features. Following greedy feature selection, feature importance was assessed using SHapley Additive exPlanations (SHAP) values (67).

### (c) Comparison of predictive performance with other *in silico* tools

To directly compare the performance of our models with other established computational tools, the blind test set was used for performance benchmarking. Additionally, our models were evaluated on an independent, ACMG/AMP-based dataset of hTERT missense variants to assess their generalizability. In the absence of hTERT*-*specific predictive tools, only conservation-based and deep-learning methods were included in our comparison.

#### Conservation-based methods

PolyPhen2 (68) and PROVEAN (69) were assessed using their recommended cutoffs of 0.45 and –2.5, respectively.

#### Deep-learning methods

AlphaMissense classifies variants with scores between 0 and 0.34 as ‘benign’, and those above 0.56 as ‘pathogenic’ (70). Variants with scores between 0.34 and 0.56 fall into an ‘ambiguous’ category and are not classified by the model. For our benchmarking, variants with scores below 0.45 were reclassified as ‘benign’, and those with scores of 0.45 or higher as ‘pathogenic’. For ESM-1b, the cutoff used was –7.5 as reported by its authors (71).

## Results

### Distribution of hTERT variants

The *TERT* manually curated dataset included 127 variants designated as pathogenic, which were distributed across the different domains. Specifically, 13% of these variants were located in the TEN domain, 17% in the RBD domain, and 21% in the CTE domain. Notably, 49% of the pathogenic variants occurred in the RT domain, showing an imbalance in their distribution (see Figure 3A and Figure 3C). On the other hand, among 59 variants designated as benign in the dataset, approximately 54% were located in the RT domain, 29% in the RBD domain, and 17% in the CTE domain. Remarkably, no benign variants were located in the TEN domain (detailed in Figure 3B and Figure 3C).

**Figure 3:**
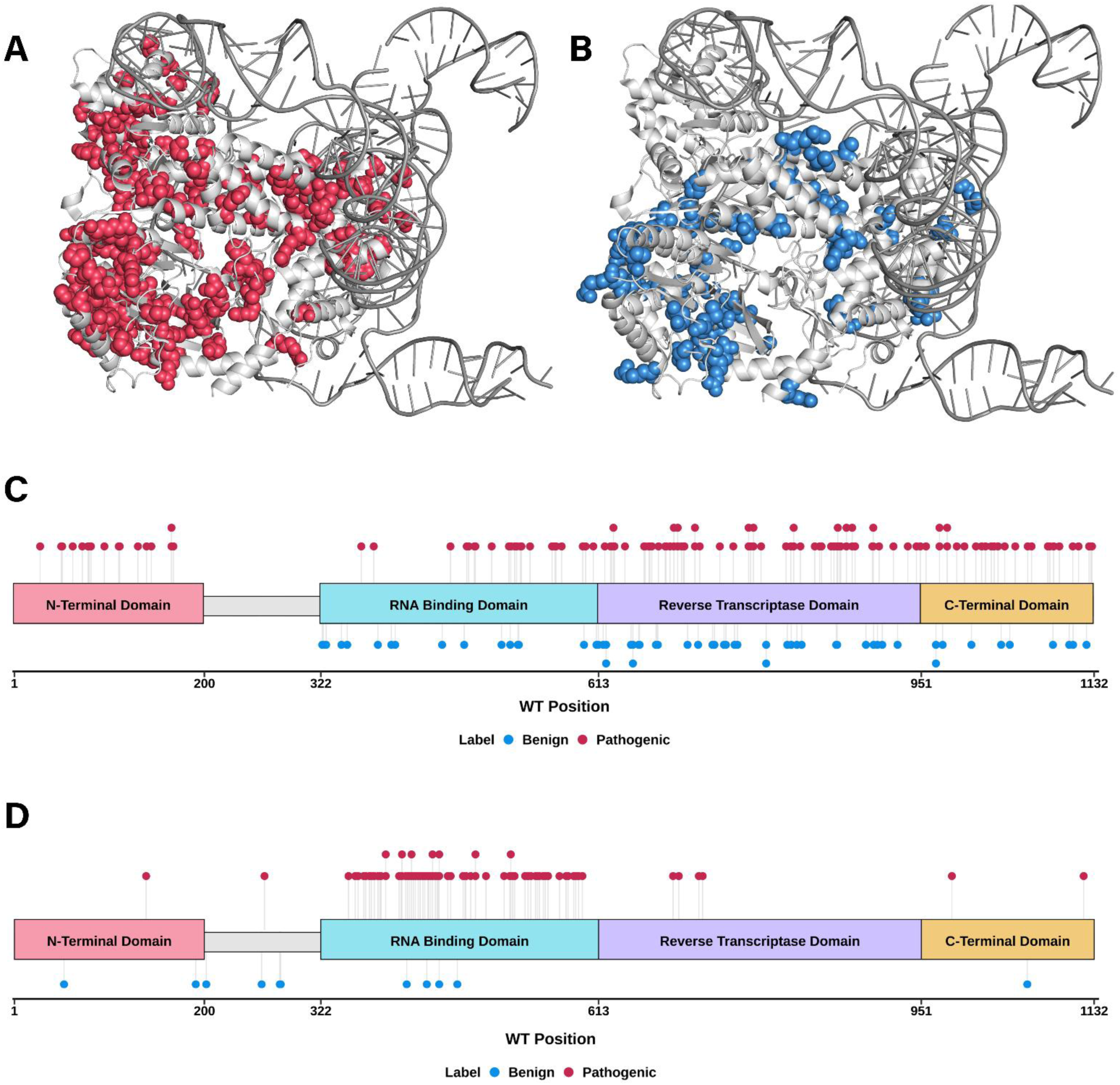
Distribution of missense variants in hTERT. Distribution of pathogenic and benign missense variants in the *TERT* manually curated dataset on the PDB structure 7QXA (13) (**A, B**) and sequence (**C**) of hTERT, as well as the distribution of pathogenic and benign variants of the ACMG/AMP-based dataset in the protein sequence (**D**). The red labels represent the pathogenic variants, while the blue ones represent the benign. The four functional domains of hTERT are represented with colours pink, cyan, purple, and yellow, respectively.

The uneven distribution of pathogenic and benign variants across the hTERT domains could be influenced by several factors, including the functional significance of each domain, the extent to which they have been characterized, and ascertainment bias. The RT domain, comprising 338 amino acids, is the largest structural region of hTERT and houses the catalytic core essential for telomerase activity.

This critical role has likely led to more extensive characterization, resulting in a higher number of variants being reported in this region. In contrast, the TEN domain, which spans 200 amino acids, lacked benign variants in the dataset, possibly reflecting a lower degree of characterization due to its less direct involvement in enzymatic activity.

This dataset comprised 135 unique positions along the hTERT sequence, with 25 positions showing multiple amino acid substitutions. Among these, 7 positions exhibited both benign and pathogenic variants, indicating variability in the pathogenicity at these sites.

Conversely, in the ACMG/AMP-based dataset, which included a total of 77 pathogenic variants, 90% were found predominantly in the RBD domain, while the RT, CTE, and TEN domains contained significantly fewer pathogenic variants (refer to Figure 3D). This distribution can be partially attributed to the fact that the RBD has previously been found to be particularly intolerant to variation (72,73), resulting in a higher likelihood that the ACMG/AMP code applying to a “mutational hotspot”. Notably, this dataset only included 11 benign variants, primarily (36%) located in the linker region between the TEN and the RBD domains and within the RBD domain itself (36%), whereas no benign variants were found in the RT domain. Moreover, the ACMG/AMP-based dataset included 7 positions with multiple variants, with only 1 showing differing classifications.

Additionally, the frequency of each type of amino acid change included in our curated dataset was analysed (Figure S2). The results revealed that amino acid substitutions involving hydrophobic or positively charged residues collectively accounted for 61% of the observed changes.

This aligned with previous findings, which highlighted the critical role of hydrophobic residues in maintaining the stability of globular proteins (74). Hydrophobic residues are essential for stabilizing the three-dimensional structure of these proteins, and disruptions in these residues often leads to protein misfolding, destabilization, or aggregation, thereby compromising both protein stability and function. Similarly, positively charged residues are essential for mediating electrostatic interactions that stabilize protein-nucleic acid complexes (75). Given that hTERT contains highly charged regions to make closer contact with its RNA subunit and the RNA–DNA heteroduplex (14, 76), it is particularly susceptible to changes in charged residues, with even minor alterations destabilizing its structure and function (77). These observations might underscore the significant impact of these critical amino acid types on hTERT’s stability and functionality, explaining their frequent involvement in disease-associated variation.

### Investigating potential pathogenic drivers of hTERT

To explore the potential features of hTERT that most frequently contribute to the development of TBDs when disrupted, an extensive statistical comparison was conducted using the Wilcoxon rank-sum test. This analysis aimed to identify features with significant differences (p-value < 0.05) between the pathogenic and benign variants in the *TERT* manually curated dataset.

Initially, a subset of features, including changes in protein stability, protein-nucleic acid interactions, and protein-protein contacts, were evaluated due to their relevance to hTERT’s functional role (see Figure 4A and Figure S3). No significant differences were found (p-value > 0.05) in protein stability changes measured by Dynamut2 (ΔΔG; Figure S3A) nor in protein-nucleic acid interactions, as measured by mCSM-NA (for both single-stranded DNA and RNA; Figures S3B and S3C). These results may be attributed to the relatively large distance between the variants in the dataset and the nucleic acids, with an average of 10.88Å, which may have contributed to reducing the sensitivity for detecting meaningful differences in protein-nucleic acid interactions between pathogenic and benign variants.

**Figure 4:**
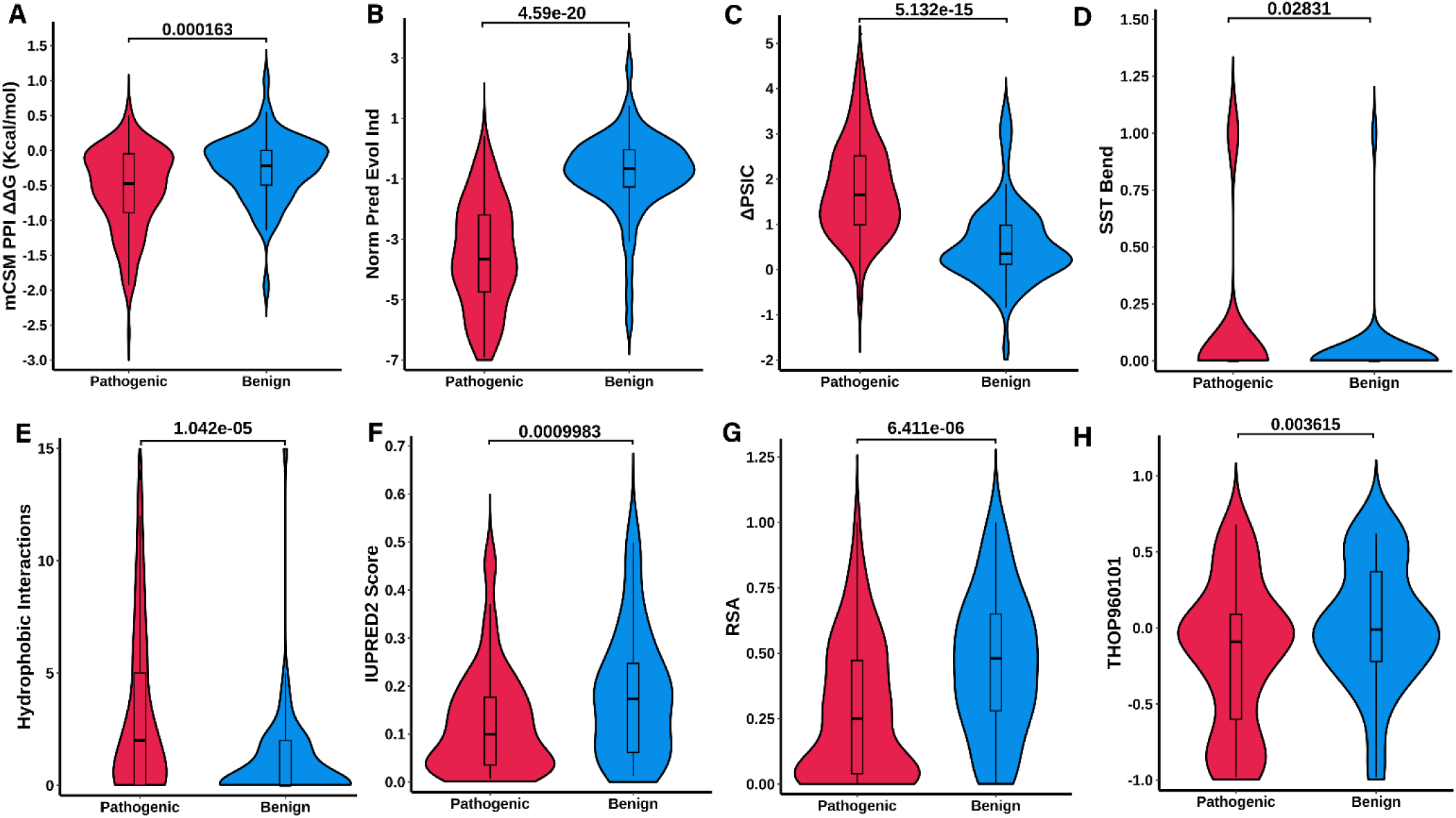
Identifying potential pathogenic drivers of hTERT. Statistical analysis was performed using the Wilcoxon rank-sum test to evaluate the variant effect on protein-protein interactions computed by mCSM-PPI (**A**); on evolutionary conservation captured by GEMME (**B**) and Envision (**C**); on biochemical properties of the mutational local environment capture by Secondary Structure Type Bend (**D**), Arpeggio Hydrophobic Interactions (**E**), IUPRED2 score (**F**), and Relative Solvent Accessibility (**G**); and on amino acidic biochemical properties captured by Aaindex (**H**). The analysis was conducted to identify significant differences between pathogenic and benign variants, with a significance level of 5%.

However, notable changes were observed in protein-protein binding affinities captured by mCSM-PPI (ΔΔG; Figure 4A). To specifically assess the impact of variants on hTERT-TPP1 interactions, H2A-H2B as well as DNA were removed from the 7QXA structure before inputting it into mCSM-PPI, ensuring that only hTERT-TPP1 interactions were captured. The average mCSM-PPI score for all variants included in the dataset was –0.463 Kcal/mol, indicating a general destabilizing effect on protein interactions. Notably, pathogenic variants had a more destabilizing average score of –0.559 Kcal/mol, while benign variants had a less pronounced destabilizing effect, with an average score of –0.245 Kcal/mol. A Wilcoxon rank-sum test performed on these scores revealed a statistically significant difference (p-value = 2.22×10^-5^) between pathogenic and benign variants. Given that TPP1 plays a crucial role in telomerase recruitment to telomeres, these findings suggest that pathogenic variants may disrupt this interaction, potentially contributing to the development of TBDs (78).

We then expanded our analysis to include 295 features, identifying 69 that exhibited significant differences (p-value < 0.05) between pathogenic and benign variants (included in **Table S1**). These features were distributed across three main categories: conservation scores, variant local environment, and biochemical properties.

### Conservation Scores

Several evolutionary conservation metrics, including GEMME, ConSurf, Position-Specific Independent Counts (PSIC) scores, and Mutation Multiple Sequence Alignment (MSA) congruency, revealed significant differences between pathogenic and benign variants (p-values ranging from 4.95×10^-20^ to 0.0137, Figure 4B, C, Figure S4 and Table S1). Population conservation scores from MTR3D, MTR v.1, and MTR v.2 further supported these findings, with significant p-values ranging from 9.62×10^-06^ to 0.0413 (Table S1), consistently highlighting the high evolutionary conservation of hTERT. Notably, GEMME ΔΔE scores demonstrated a significant distinction between variant types, with pathogenic variants averaging –3.2 compared to –0.86 for benign variants (Figure 4B). Variants with ΔΔE values closer to 0 are indicative of tolerant substitutions, while those closer to -6 correspond to non-tolerated changes (57). Therefore, these findings suggested that pathogenic variants tend to occur at highly conserved residues, where alterations are less tolerated and more likely to disrupt protein function. Additionally, delta PSIC (ΔPSIC), which measures the conservation difference between wild-type and mutant residues, also showed significant differences, with pathogenic variants having a mean ΔPSIC of 1.8 versus 0.567 for benign variants (Figure 4C). These results further suggested that pathogenic variants occur at more conserved positions, which are often important to maintain protein function.

#### Local Environment

Biophysical measurements, including the Relative Solvent Accessibility (RSA), Contact Area (CA) Depth, Relative B factor, Secondary Structure Types (SST), IUPRED2, IUPRED3, Anchor, and interactions analysed using Arpeggio were significantly different between pathogenic and benign variants (p-values between 6.41×10^-06^ and 0.0283, as shown in Figure 4D-G, Figure S5 and Table S1). Pathogenic variants tended to localize in protein bends (Figure 4D), which were particularly influential in distinguishing variant types. Moreover, among the interactions captured by Arpeggio, hydrophobic, proximal, carbon-protein, and weak polar interactions were highlighted in the analysis. For instance, pathogenic variants exhibited an average of 4 hydrophobic interactions, while benign variants had only 1 (Figure 4E), suggesting that pathogenic variants occurred at residues essential for maintaining protein stability. These residues are often located in the protein core, where changes in hydrophobic contacts can significantly destabilize hTERT’s native structure. In contrast, benign variants maintained a smaller difference in hydrophobic contacts, indicating a lower impact on overall protein stability and function.

Notably, IUPRED2 scores, which measure protein intrinsic disorder, revealed that pathogenic variants had an average score of 0.125, significantly lower than the mean value of 0.182 for benign variants (Figure 4F). This indicated that pathogenic variants are more likely to occur in structured regions, potentially contributing to their disruptive effects on protein function.

Pathogenic variants had an average RSA of 0.277, compared to 0.457 for benign variants (Figure 4G). This significantly lower RSA for pathogenic variants suggested that they are more likely to occur in buried regions of the protein, which are crucial for maintaining structural stability (79). Such variants may disrupt the protein’s core integrity, leading to potential functional impairments and disease. In contrast, benign variants, which tend to occur in more exposed regions, are less likely to impact protein function.

#### Biochemical Properties

Significant biochemical features included several Aaindex measurements, PAM and BLOSUM scores, and amino acid polarity (p-values ranging from 0.002 to 0.0451, depicted in Figure S6 and Table S1).

Additionally, THOP960101 Aaindex score, which reflects protein contact potentials, showed that pathogenic variants had a mean score of –0.246, while benign variants had an average of 0 (Figure 4H). These findings highlighted that pathogenic variants are associated with less favourable or destabilizing interactions, which may disrupt protein stability and function. In contrast, benign variants tend to have neutral interactions, suggesting that they are less likely to impact protein stability, implying a lower risk of disrupting the protein’s structural integrity.

### Predicting hTERT variant pathogenicity with machine learning

Building on these findings, we employed a machine learning-based approach to further elucidate the relationship between hTERT variants and pathogenicity. We developed three different models, all based on the Random Forest algorithm, named Position 10CV, Domain 4CV, and Domain 3CV.

The primary distinction between these models lay in their cross-validation strategies. The Position 10CV grouped variants into different folds based on their specific residue positions within the hTERT protein sequence, using a stratified 10-fold cross-validation strategy. In contrast, the Domain 4CV and Domain 3CV models grouped variants by the domains they were located in. Due to structural constraints in the PDB structure *7QXA*, which lacked the linker region between the TEN and RBD domains, only the four functional domains were considered, making 10-fold cross-validation infeasible. Consequently, the Domain 4CV model employed a 4-fold cross-validation approach, with each fold consisting of variants from a single domain, while the Domain 3CV model employed a 3-fold cross-validation approach.

Model performance was evaluated using the MCC metric on a blind test set (see Figure 2; results in Table 1) and an independent dataset curated following ACMG/AMP guidelines (Table 2). This additional validation provided insights into the models’ performance on rigorously classified variants, thereby assessing its applicability in both a research and clinical context. On the blind test set, both the Position 10CV and the Domain 3CV models achieved the highest performance (MCC = 0.88), whereas on the independent dataset, the Position 10CV (MCC = 0.47) outperformed the domain-based models.

**Table 1:** Predictive performance of different variant effect predictors on the curated dataset. Predictive performance is assessed by Matthew’s correlation coefficient (MCC), accuracy, sensitivity, and specificity.

| Models | Method Type | Set | MCC | Accuracy | Sensitivity | Specificity |
| --- | --- | --- | --- | --- | --- | --- |
| Position 10CV | *hTERT specific* | 10-CV | 0.68 | 0.86 | 0.91 | 0.76 |
| Domain 3CV |  | 10-CV | 0.69 | 0.87 | 0.91 | 0.78 |
| Domain 4CV |  | 10-CV | 0.67 | 0.86 | 0.89 | 0.78 |
| Position 10CV |  | Blind Test | 0.88 | 0.95 | 0.83 | 1 |
| Domain 3CV |  | Blind Test | 0.88 | 0.95 | 1 | 0.83 |
| Domain 4CV |  | Blind Test | 0.76 | 0.90 | 0.92 | 0.83 |
| AlphaMissense | *Deep learning-based* | Blind Test | 0.29 | 0.47 | 0.23 | 1 |
| ESM-1b |  | Blind Test | 0.31 | 0.47 | 1 | 0.29 |
| PolyPhen2 | *Conservation-Based* | Blind Test | 0.76 | 0.90 | 0.92 | 0.83 |
| PROVEAN |  | Blind Test | 0.65 | 0.79 | 0.69 | 1 |

**Table 2:** Predictive performance of variant effect predictors on the ACMG/AMP-based dataset. Predictive performance is assessed by Matthew’s correlation coefficient (MCC), accuracy, sensitivity, and specificity.

| Models | Method Type | MCC | Accuracy | Sensitivity | Specificity |
| --- | --- | --- | --- | --- | --- |
| Position 10CV | *hTERT- specific* | 0.47 | 0.77 | 0.75 | 0.91 |
| Domain 3CV |  | 0.37 | 0.68 | 0.65 | 0.91 |
| Domain 4CV |  | 0.37 | 0.77 | 0.78 | 0.73 |
| AlphaMissense | *Deep learning-based* | 0.14 | 0.25 | 0.14 | 1 |
| ESM-1b |  | 0.29 | 0.5 | 0.43 | 1 |
| PolyPhen2 | *Conservation-Based* | 0.17 | 0.65 | 0.66 | 0.58 |
| PROVEAN |  | 0.24 | 0.41 | 0.33 | 1 |

These results highlighted critical differences in the models’ ability to generalize. The Position 10CV, by grouping variants at the residue level, effectively captured subtle pathogenic patterns across diverse variants, which translated into superior generalization to clinical data. In contrast, the Domain 4CV and Domain 3CV models achieved higher performance on the blind test set but were less effective on the independent dataset, indicating potential overfitting to domain-specific features that reduced their generalizability to clinical scenarios, and therefore reducing their clinical applicability.

The Domain 4CV model, employing a Leave-One-Group-Out cross-validation strategy by being trained on variants from three domains and tested on the remaining one during the k-fold cross-validation, provided a closer examination of domain-specific performance (see Table 2). For example, when trained on variants from the RBD, RT, and CTE domains and tested on the TEN domain, it achieved an MCC score of 0 due to the absence of benign variants in the TEN domain. Similarly, the model exhibited the lowest performance on the RBD domain (MCC = 0.47), likely reflecting the inherent complexity of this domain and insufficient feature representation for benign variants in this region. These results underscored the impact of data imbalance in the original dataset, where pathogenic and benign variants were unevenly distributed across domains. Such imbalance likely biased the feature learning process, particularly in the RT domain, and hindered the domain-based model’s ability to generalize to clinical data.

Despite these challenges, our models demonstrated high predictive performance across datasets. However, they faced challenges in accurately classifying some hTERT variants (see Table S3 and Table S4). Substitutions involving proline residues (e.g., P785L and P703L) were particularly difficult for the models, although they represented around 13% of the variants included in the *TERT* manually curated dataset. Proline plays a critical role in protein folding and stability, a property that likely contributes to the complexity of accurately predicting the impact of such variants (80). Additionally, variants involving hydrophobic (e.g., A670V and A67V) and polar (e.g., N906S) residues posed further classification challenges due to their complex biochemical roles in protein structure and function. It is important to note that these “misclassified” variants may not necessarily be misclassified, as there is inherent uncertainty around the accuracy of their assigned labels. Refining our models to address these complexities will ultimately improve the accuracy of pathogenicity predictions, particularly in structurally and functionally important areas.

### Performance comparison with state-of-the-art methods

When compared to existing *in silico* tools, our models consistently outperformed them across both the blind test set and the ACMG/AMP-based dataset, demonstrating their superior predictive capabilities across several evaluation metrics (see Table 1 and Table 2).

In the blind test set, the Position 10CV (MCC = 0.88), Domain 3CV (MCC = 0.88), and Domain 4CV (MCC = 0.76) models outperformed the state-of-the-art methods, including the top-performing benchmark tool, PolyPhen2 (MCC = 0.76). These results underscored the enhanced ability of our models to incorporate key features related to pathogenicity in hTERT variants, thus demonstrating their superior predictive accuracy.

Similarly, our models exhibited superior performance on the ACMG/AMP-based dataset. The Position 10CV (MCC = 0.47; accuracy = 0.77; sensitivity = 0.75) and Domain 4CV (MCC = 0.37; accuracy = 0.77; sensitivity = 0.78) models significantly outperformed ESM-1b (MCC = 0.29; accuracy = 0.5; sensitivity = 0.43), the top-performing state-of-the-art method for this dataset. This reinforced the robustness of our models in classifying clinical variants with high sensitivity and accuracy.

Moreover, our models demonstrated superior generalization capabilities as evidenced by the Area Under the Curve (AUC) scores. The Position 10CV (AUC = 0.870) and Domain 4CV (AUC = 0.821) models achieved notably higher AUCs compared to AlphaMissense (AUC = 0.808), as illustrated in Figure 5A and Figure 5B. These results further validated the clinical relevance and predictive accuracy of our models in distinguishing pathogenic from benign variants in an ACMG/AMP-based dataset.

**Figure 5:**
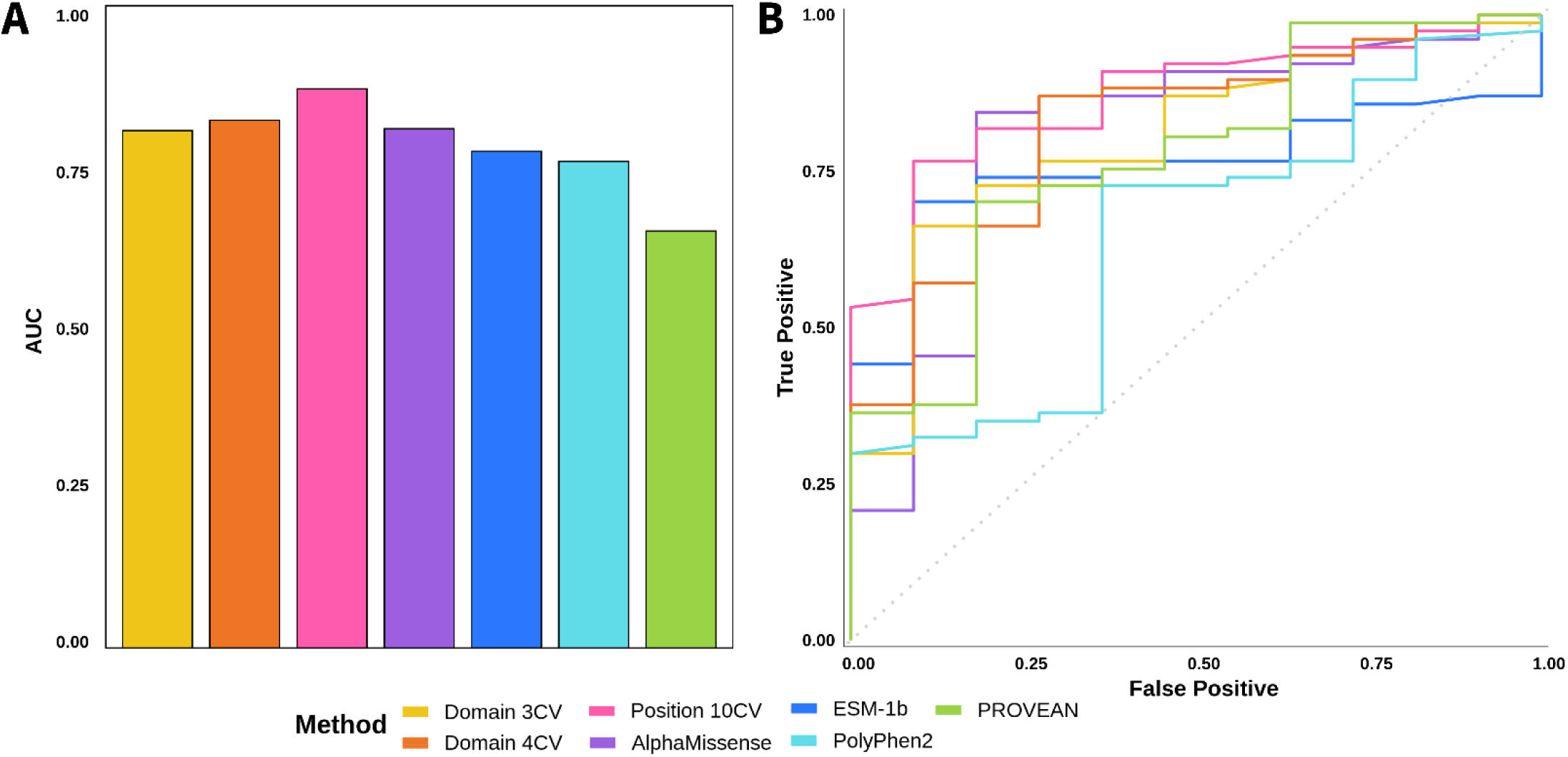
Predictive performance comparison. Predictive performance of the Position 10CV, Domain 3CV, and Domain 4CV models compared to state-of-the-art methods using the ACMG/AMP-based dataset. Performance is assessed by AUC scores (**A**) and the receiver operating characteristic (ROC) curves (**B**).

While PolyPhen2 exhibited comparable performance in the blind test set (MCC = 0.76), it is important to note that its training datasets included 20 hTERT variants, 17 of which overlapped with the *TERT* manually curated dataset and 2 with the ACMG/AMP-based dataset (68). Similarly, PROVEAN was trained on 35 hTERT variants, 28 of which overlapped with the *TERT* manually curated dataset, and 4 with the ACMG/AMP-based dataset (69). These overlaps likely introduced bias in the performance comparisons. In contrast, our blind test set and the ACMG/AMP-based dataset did not include any variants occurring at the same positions as those in our training and validation sets, ensuring a more rigorous and unbiased evaluation.

Among all models, the Position 10CV model consistently demonstrated the highest generalization capability and strongest overall performance on the ACMG/AMP-based dataset across several metrics. Therefore, this model represented the most robust and reliable method for predicting the pathogenic effects of hTERT variants, marking a significant advancement over exiting *in silico* tools.

### Interpreting features in hTERT variants

Since both Position 10CV and Domain 4CV demonstrated strong performance in predicting the pathogenic effects of hTERT missense variants, we applied feature interpretation methods to assess the contribution of each feature to the models’ predictions. This analysis provided valuable insights into potential pathogenic drivers of hTERT, enhancing our understanding of the underlying mechanisms. Conversely, the Domain 3CV model, which demonstrated a significant performance deterioration on the ACMG/AMP-based dataset, was excluded from further analysis due to its reduced generalization capability, limiting its reliability in exploring potential pathogenic drivers of hTERT.

#### Position 10CV

The most contributing feature for this model was the normalized evolutionary combined score (normPred_evolCombi), obtained from GEMME, which evaluated the impact of missense variants based on evolutionary conservation. Low values of this score were associated with pathogenic variants (see Figure S3B and Figure 6A), indicating that variants in highly conserved regions are more likely to disrupt hTERT function and structure, leading to a pathogenic phenotype.

**Figure 6.**
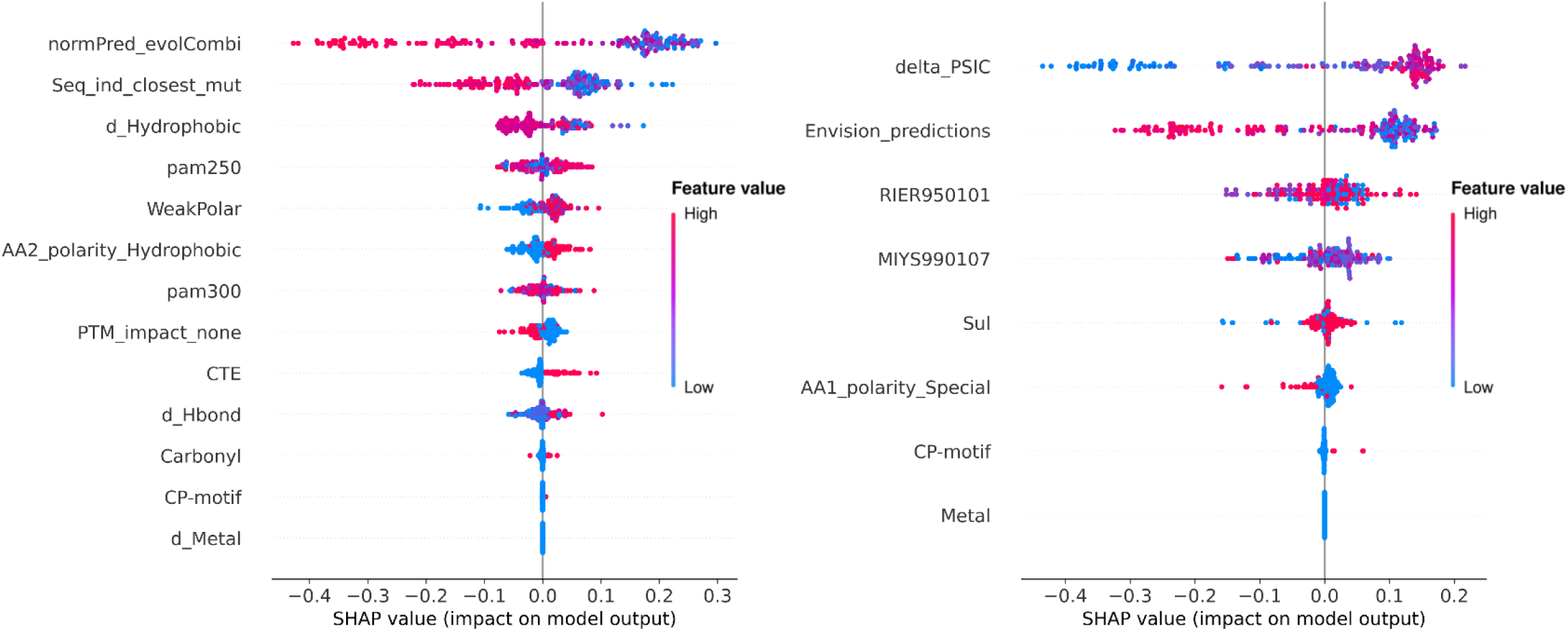
Model interpretation using SHAP values. Feature importance of the Position 10CV (**A**) and Domain 4CV (**B**) models evaluated using SHapley Additive exPlanations (SHAP). Features were ordered from top to bottom with respect to their contributions to the models. The contribution of the different features towards pathogenicity is represented by the x-axis (positive values: pathogenic, negative values: benign), while the value of each feature is represented by color (red: high values of the feature, blue: low values of the feature). Combining both aspects, the contribution of each feature to the model can be extracted. Highlighted features include GEMME (normPred_evolCombi); Envision (seq_ind_closest_mut, AA2_polarity_Hydrophobic, delta_PSIC, Envision_predictions, AA1_polarity_Special); Aaindex, (RIER950101, MIYS990107); Domain information (CTE, CP-motif); PAM matrix (pam250, pam300); Arpeggio (d_Hydrophobic, WeakPolar, d_Hbond, Carbonyl, Metal, d_Metal); ActiveDriver DB (PTM_impact_none); and Graph-based signatures (Sul).

The second most significant feature was the sequence-based identity score of the closest homolog with a different residue (Seq_ind_closest_mut), extracted from Envision. This feature measured how similar the mutated sequence was to the most closely related homologous sequence where the wild-type amino acid is absent. Similarly to the evolutionary score, low values of this features were associated with pathogenic variants, suggesting that these variants are more likely to occur at highly conserved positions, where sequence deviations are rare.

Finally, the third most important feature was the changes in the number of hydrophobic interactions (d_Hydrophobic), collected from Arpeggio. Hydrophobic interactions play a crucial role in protein folding and stability (81–83). In our model, a reduction in these interactions was associated with pathogenic variants, suggesting that such changes may adversely affect protein conformation and functionality.

#### Domain 4CV

The most contributing feature was the difference in Position-Specific Independent Counts (ΔPSIC), obtained from Envision, which measured the likelihood of each amino acid occurring at a specific position in the protein sequence based on evolutionary information. High values of this feature were associated with pathogenic variants (see Figure 6B), suggesting that these changes were less tolerated and more likely to disrupt protein function. The second most contributing feature was Envision predictions, which provides a quantitative measure of the impact of missense variants on protein function. Lower scores were associated with higher pathogenicity, indicating pathogenic variants are likely to disrupt protein function due to significant effects on structure, stability, or conserved functional regions.

Finally, the third most contributing feature was RIER95010, collected from Aaindex, which measures the stabilization parameter for alpha-helices (ΔΔG). Lower values of this feature were associated with pathogenic variants, indicating a greater destabilizing effect on alpha-helices. This suggests that these variants may compromise the structural integrity of the protein, potentially impacting its overall function.

These findings underscored the limitations of relying exclusively on sequence-based features for accurately predicting pathogenic variants in hTERT. Traditional *in silico* tools, which depend exclusively on sequence-based features, significantly underperformed compared to our Position 10CV and Domain 4CV models, demonstrating their limitations and underscoring the need for more comprehensive approaches.

While the three models outperformed state-of-the-art methods by incorporating structural information, the Position 10CV model demonstrated superior generalization on the ACMG/AMP-based dataset, as well as the best overall performance on the blind test set. The advantage of the Position 10CV model likely lie in its broader set of structural features, which enabled it to effectively capture the structural changes induced by missense variants. Furthermore, this model employed a more effective representation of the data, minimising the risk of data leakage, which refers to the unintentional inclusion of information from the test set during the training phase, thereby inflating model performance. In contrast, domain-specific splits used in the Domain 4CV and Domain 3CV models were more prone to data leakage due to insufficient data in certain domains. The Position 10CV model’s robust handling of data distribution, combined with its ability to integrate diverse structural insights, contributed to its superior predictive performance on the ACMG/AMP-based set.

### Exploring hTERT mutational landscape through computational saturation mutagenesis

Given the superior and consistent performance of our Position 10CV model, particularly in the ACMG/AMP-based dataset, the comprehensive mutational landscape of hTERT was calculated (Figure 7) based on predictions from this model. The prediction for all possible missense variants is shown in Figure 7D, while the average pathogenicity across different positions is illustrated in Figure 7C. To enhance the interpretation of these results in the context of hTERT’s structure, we mapped the average pathogenicity predictions onto the 7QXA PDB structure (Figure 7A) and compared them with AlphaMissense predictions (Figure 7B). In contrast to AlphaMissense, which classified many residues as ‘ambiguous’, our model provided clearer predictions, categorising variants as either pathogenic or benign.

**Figure 7:**
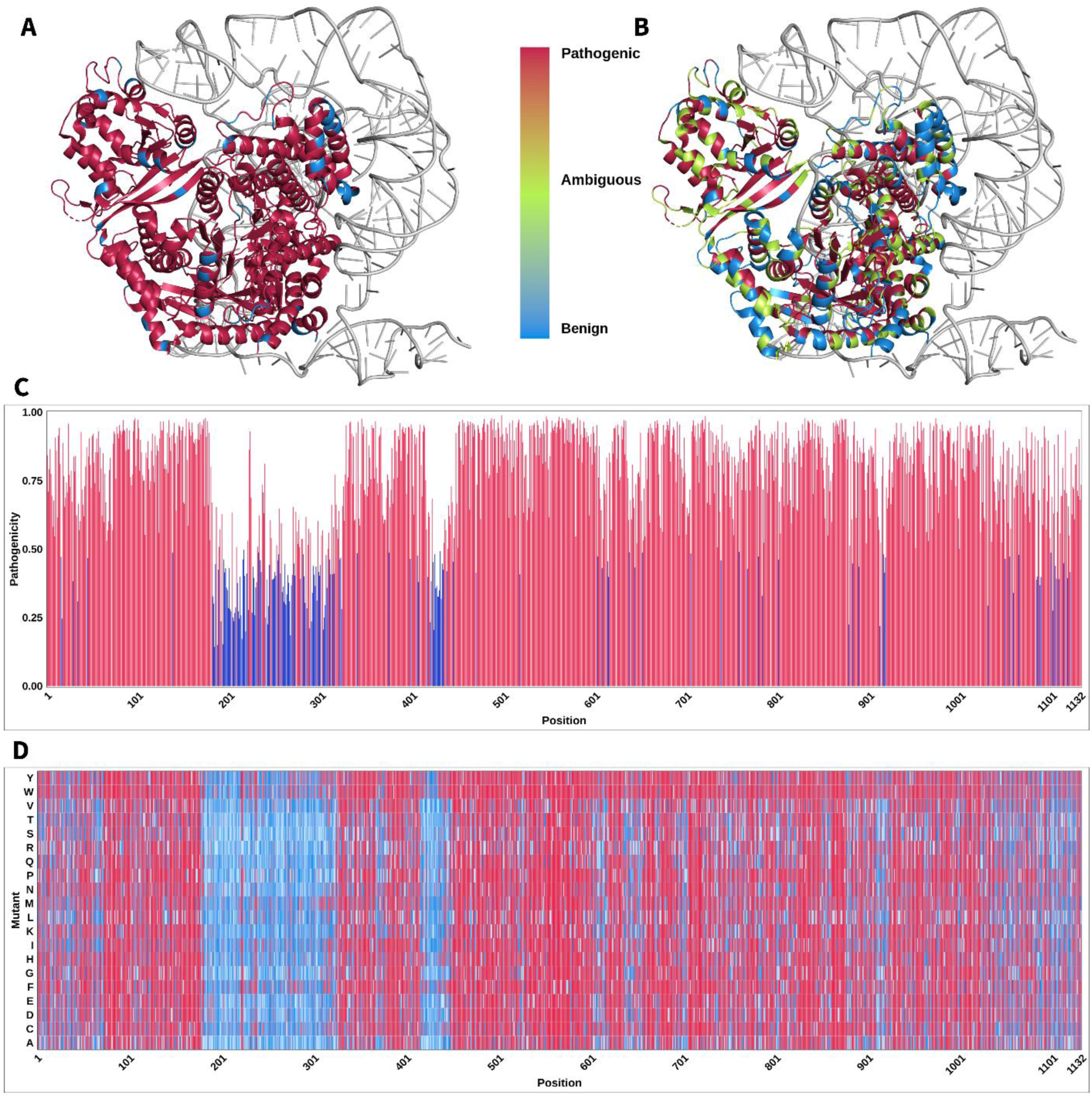
Computational saturation mutagenesis analysis. The pathogenicity of hTERT was predicted using the model and visualized as follows: (**A**) The average pathogenicity across the hTERT sequence, mapped onto the 7QXA PDB structure; (**B**) AlphaMissense predictions, overlaid on the 3D structure; (**C**) Average pathogenicity shown on the hTERT amino acid sequence; (**D**) Detailed examination of mutant-specific pathogenicity.

When analysing the 7QXA structure, we noted that due to loop flexibility, several regions were missing residues, specifically hTERT residues 1-3, 105-111, 180-321, and 418-443. Since understanding the effect of variant in a structural context is essential, we used the experimental 7QXA structure to map those variants that fall within its resolved regions. However, to account for the missing regions caused by loop flexibility, a previously published homology model based on another structure (PDB ID: 7BG9) was used as a reliable alternative for calculating structural features for variants located within the missing regions of the 7QXA structure (84). The 7BG9 structure was chosen due to its high structural similarity with 7QXA (RMSD = 1.104 Å across 10,643 aligned atoms).

Despite the absence of experimental density for these regions in 7QXA, our Position 10CV model accurately predicted 24 of the 32 benign variants available in ClinVar for these regions, highlighting its predictive capacity. Interestingly, substitutions of several residues in the linker region between the telomerase essential N-terminal Extension (TEN) domain and the RNA Binding Domain (RBD) were predicted to be pathogenic (Table S5).

Notably, nine residues within this linker, predicted to be pathogenic when changed, belong to the bipartite nuclear localization signal (NLS), spanning amino acids 222-240. This NLS is crucial for hTERT’s nuclear import (85), and variants in these residues may significantly disrupt nuclear localization, emphasizing their functional importance. Specifically, phosphorylation at the Serine residue 227 (S227) is essential for hTERT’s nuclear translocation. Remarkably, our Position 10CV model identified 6 variants at S227 (S227F, S227I, S227M, S227V, S227W, and S227Y) as pathogenic. These variants, characterized by changes in size and hydrophobicity, likely disrupt phosphorylation, impairing nuclear transport of hTERT. Remarkably, these variants were not captured by AlphaMissense or ESM-1b, while PROVEAN only identified S227F and S227W as pathogenic. This highlights the strength of our model in detecting pathogenic variants at critical functional residues of hTERT.

Moreover, tryptophan (Trp) exhibited the highest average pathogenicity among all residues, closely followed by tyrosine (Tyr) and phenylalanine (Phe) (see Figure S7). These three residues accounted for approximately 15% of the variants in the training dataset, appearing in both wild-type and mutant forms. Notably, every variant involving Trp, Tyr, or Phe in the training set was classified as pathogenic, which likely contributed to their high pathogenicity scores in our model. In contrast, while glutamate (Glu) comprised 6 out of 7 variants classified as pathogenic in the training set, it exhibited a significantly lower average pathogenicity overall. This distinction highlighted the robustness of our model in discerning not only the presence of pathogenic variants but also their varying degrees of impact, further validating its predictive capability.

The unique structural properties of Trp, Tyr, and Phe, particularly their aromatic rings and bulky side chains, likely play a critical role in this outcome. These residues are often found in the protein core, where their size and hydrophobic characteristics can induce significant changes in the local protein conformation, disrupting existing interactions and leading to potential loss of function (86,87).

### Validation of position 10CV model predictions against experimental literature

To further validate the predictive capability of the Position 10CV model, we conducted an independent evaluation using experimentally characterized hTERT variants that all have robust published telomerase functional data available and are not included in the previous datasets (Tables S6-S9). This external validation aimed to assess the generalization of the model beyond the training data and to benchmark its performance against state-of-the-art methods. By evaluating our model’s predictions on variants with known effects on telomerase function, we aimed to mitigate potential biases and overfitting, reinforcing the robustness and reliability of our approach.

The Position 10CV model exhibited superior predictive performance (MCC = 0.49), outperforming conservation-based and deep learning-based approaches in classifying hTERT variants (Table 3). Notably, deep learning models such as AlphaMissense (MCC = 0.27) and ESM-1b (MCC = 0.39) exhibited lower performance, likely due to their reliance on broad evolutionary and structural features without being tailored to hTERT.

**Table 3.**
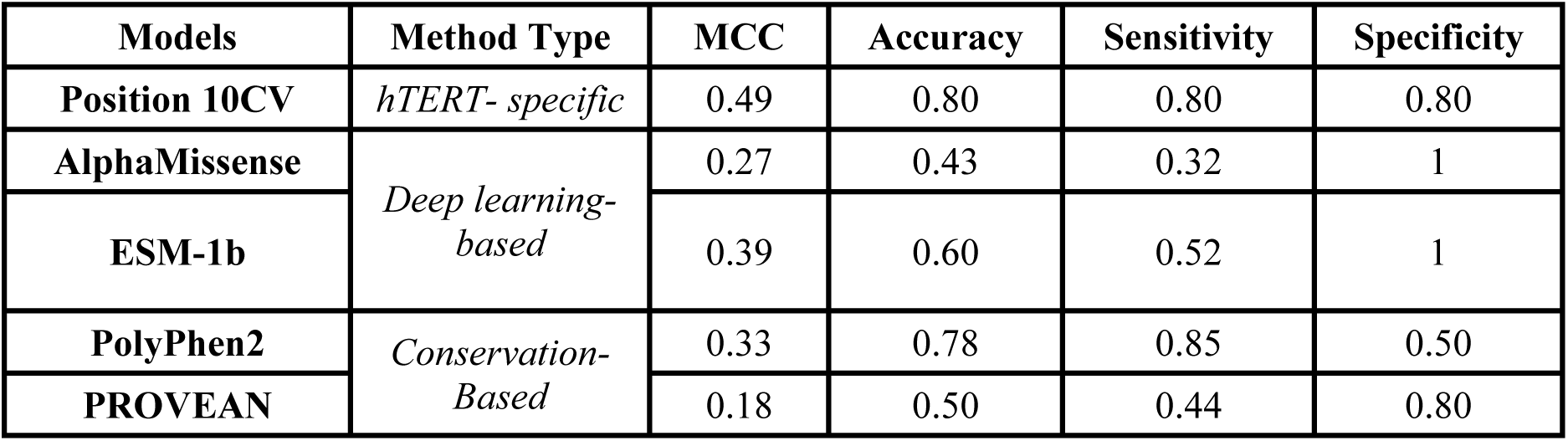
Predictive performance of variant effect predictors on experimental literature. Predictive performance is assessed by Matthew’s correlation coefficient (MCC), accuracy, sensitivity, and specificity.

| Models | Method Type | MCC | Accuracy | Sensitivity | Specificity |
| --- | --- | --- | --- | --- | --- |
| **Position 10CV** | *hTERT- specific* | 0.49 | 0.80 | 0.80 | 0.80 |
| **AlphaMissense** | *Deep learning-based* | 0.27 | 0.43 | 0.32 | 1 |
| **ESM-1b** |  | 0.39 | 0.60 | 0.52 | 1 |
| **PolyPhen2** | *Conservation-Based* | 0.33 | 0.78 | 0.85 | 0.50 |
| **PROVEAN** |  | 0.18 | 0.50 | 0.44 | 0.80 |

Among conservation-based methods, PolyPhen2 achieved relatively high sensitivity (0.85) but lower specificity (0.5), indicating a tendency to overpredict pathogenic variants. Similarly, PROVEAN exhibited low overall performance (MCC = 0.18, accuracy = 0.50), reinforcing the limitations of generic conservation-based approaches in capturing the nuances of hTERT. In contrast, our Position 10CV model balanced both sensitivity (0.80) and specificity (0.80), providing a more accurate classification of pathogenic and benign variants.

These findings underscore the advantages of a model specifically tailored to hTERT. By integrating structural, functional, and evolutionary features tailored to hTERT, the Position 10CV model achieved more reliable predictions, surpassing conventional *in silico* predictors in an independent experimental dataset. This highlights its potential applicability in variant interpretation, providing valuable insights that could inform clinical decision-making and drive future research into telomere biology disorders.

### CharacTERT webserver

To furthas been developed as a user-friendly, interactive webserver that allows researchers to explore the pathogenicity effects of all possible variants in hTERT. The webserver is freely accessible to the research community at https://biosig.lab.uq.edu.au/charactert.

By selecting a specific variant of interest, users can view detailed structural information at the variant site, including metrics such as the distance to the protein-nucleic acid interface, relative solvent accessibility, and 3D visualisation of interatomic interactions in both wild-type and mutant structures (88). Furthermore, users can download the saturation mutagenesis predictions generated by our Position 10CV model for further analysis and application.

## Discussion

The catalytic component of the human telomerase holoenzyme, telomerase reverse transcriptase (hTERT), has been a focus of study due to its critical role in cancer, cellular aging, and Telomere Biology Disorders. Missense mutations in hTERT are a common cause of inherited telomere disorders as they can significantly alter protein structure and disrupt its normal function. Given the extensive time and resources required, experimentally elucidating the phenotype of all possible hTERT variants is impractical. To address this challenge, we conducted a comprehensive computational analysis and developed three models that integrate sequence and structural features to facilitate the characterization of the pathogenic effects of missense variants in hTERT, thereby enhancing predictive performance.

Our analysis identified several potential pathogenic drivers through statistical comparisons and subsequent machine learning. These encompassed a wide range of biological features, including evolutionary and population conservation (e.g., GEMME, MTR scores), protein stability and structural impact (e.g., PSIC, RSA, mCSM), changes in protein-protein interactions (e.g., van der Waals clashes, Weak Polar interactions), altered protein disorder and flexibility (e.g., IUPRED scores, Bend Secondary Structure Type), and key amino acid properties (e.g., wild-type hydrophobicity, Aaindex scores).

The structural complexity and functional significance of specific domains, particularly the RNA Binding and Reverse Transcriptase domains, posed challenges for accurate variant classification. Across our models, approximately 70% of misclassified variants were located within these two domains. The intricate structures and dynamic conformational changes associated with hTERT’s nucleic acid interactions likely contributed to a higher number of misclassified variants. For example, the RBD undergoes significant conformational shifts to accommodate RNA binding, while the RT domain participates in multiple catalytic and binding processes (89). This structural flexibility and multifunctionality introduced variability that challenged our models’ predictions. Future determination of hTERT structures at various stages of the catalytic cycle could offer valuable insights, enhancing the accuracy of variant classification in subsequent versions of our models.

Notably, evolutionary conservation was the most important feature for our models, despite leading to some misclassifications. However, it is important to note that these variants may not necessarily be misclassified, as there is inherent uncertainty in the accuracy of their assigned labels. ΔΔE scores measure the impact of variants on protein stability and function, with values near –6 indicating non-tolerated changes and values closer to 0 suggesting tolerant substitutions. Among the misclassified benign variants, the average ΔΔE score was -2.57, whereas all misclassified pathogenic variants exhibited ΔΔE values averaging -0.62 (Table S3). Therefore, refining our models to address these complexities will ultimately improve the accuracy of pathogenicity predictions, particularly in structurally and functionally important areas.

Despite these challenges, the Position 10CV model successfully classified both experimentally characterized and clinically important (ACMG/AMP) variants, demonstrating high confidence scores and consistent alignment with findings from the literature. This result underscores the reliability of our predictions and supports the fact that the model effectively captures the functional implications of hTERT variants. By correlating our findings with existing experimental research, we reinforce the validity of our approach and provide valuable insights for future research into hTERT-related pathologies.

The superior performance of our Position 10CV model can be attributed to the key features it incorporated. Feature interpretation techniques revealed that conservation-related features were among the most significant contributors, highlighting hTERT’s high degree of conservation (90,91). Variants in conserved regions are more likely to impair protein function, consistent with previous studies (92,93). Hydrophobic interactions were also crucial, aligning with their established role in mediating the RNA-binding activity of hTERT through the RBD domain (94). Similarly, weak polar interactions, another key feature in our model, have been reported to play a critical role in telomere protein binding to DNA (95).

Moreover, the Position 10CV model demonstrated superior generalization by effectively handling data distribution challenges and minimizing the risk of data leakage. Unlike the domain-specific splits used in the Domain 4CV and Domain 3CV models, which were prone to data leakage due to insufficient variants included for certain domains, our Position 10CV model employed a more robust approach that ensured comprehensive data representation, enhancing its predictive reliability.

Interestingly, the C-terminal Extension (CTE) domain was also a contributing feature in our Position 10CV model (Figure 6A). Variants located within this domain were associated with pathogenic phenotypes, while those outside it were classified as benign. The CTE domain is known to participate in numerous protein-protein interactions and plays a key role in regulating hTERT localization and processivity (96). Therefore, the classification of CTE variants as pathogenic was likely influenced by their combination with other features, such as hydrophobicity (d_hydrophobic), changes in weak polar interactions (weakpolar), and sequence conservation scores (normPred_evolCombi). This suggests that the CTE domain could be particularly sensitive to changes in hydrophobic or polar amino acids due to its role in mediating essential interactions. Disruptions in these contacts within the CTE domain could potentially impair telomerase function or its localization.

When analysing the most pathogenic mutants predicted through saturation mutagenesis, aromatic residues were highlighted. The hydrophobic nature of Trp, Tyr, and Phe, along with their ability to engage in π-π stacking interactions with other aromatic residues, can destabilize protein structure and interactions. Moreover, these residues appear to be crucial in the DNA-binding process, contributing to a pathogenic phenotype (97). These findings not only validate the robustness of our model, but also align closely with known hTERT biological mechanisms, reinforcing its predictive capacity.

While both our models and conservation-based state-of-the-art methods could accurately predict ClinVar and gnomAD variants, the currently available *in silico* tools showed significant performance deterioration when classifying a dataset manually curated using ACMG/AMP guidelines as well as variants from the experimental literature. This decline may be attributed to the limited representation of structural features in these tools. In contrast, our Position 10CV model, despite not being explicitly trained on these sets of variants, incorporated structural information that effectively captured the biological and clinical relevance of the variants and, therefore, maintained consistent performance across Clinvar-curated, ACMG/AMP-curated, and functionally characterized variants.

However, the integration of structural features allowed Position 10CV to not only maintain consistent performance across datasets, but also to offer unique insights into hTERT critical regions, especially the intrinsically disordered linker between the TEN and RBD domains. Previous studies have shown that intrinsically disordered regions (IDRs) can play critical roles in biological processes, such as signalling and regulation (98,99). Moreover, analysis of the human hTERT linker region revealed its atypical amino acid composition, with high contents of Proline (18%), Arginine (14%), and Glycine (12%) (100).

SEG analysis further identified two segments within the linker (residues 213-248 and 313-323) as low complexity (101,102), suggesting their propensity to mediate protein-protein interactions. While such low-complexity regions potentially enhance self-association, under overexpression conditions they can disrupt normal cellular functions (103,104). Therefore, the pathogenicity predictions predicted by our Position 10CV model within this linker region underscores a potential critical role of these residues in maintaining hTERT’s functional dynamics.

Additionally, the nuclear localization signal (NLS), also located within the linker region between the TEN and RBD domains, underscores the functional importance of this region. Notably, our Position 10CV model successfully identified 6 variants at S227 as pathogenic, which were not captured by state-of-the-art methods. This highlights the ability of our model to detect pathogenic variants at key functional residues of hTERT.

In summary, our models accurately characterized hTERT missense variants by integrating both structural and sequence-based information, offering a robust framework for understanding the pathogenic effects of these variants. The pathogenicity predictions from our best-performing model provide a comprehensive mutational landscape of hTERT, offering valuable insights that not only can advance research on hTERT’s role in TBDs, but also hold significant potential to aid early diagnosis and facilitate the development of personalized treatment strategies.

## Data availability

The curated CharacTERT training, validation, and testing datasets including missense variants in hTERT are available upon request.

## Supplementary data

Supplementary Data are available at Biorxiv online.

## Author contribution statement

**Georgina Becerra Parra:** Data curation; methodology; formal analysis; writing – original draft; visualization; validation. **Qisheng Pan:** Software; writing – review and editing. **YooChan Myung:** Methodology; writing – review and editing. **Stephanie Portelli:** Methodology; writing – review and editing. **Niles E. Nelson:** Data curation; writing – review and editing. **Joanne L. Dickinson:** Data curation; writing – review and editing. **Sionne E.M. Lucas:** Data curation; writing – review and editing; **Jessica K. Holien:** Methodology; formal analysis; writing – review and editing. **Tracy M. Bryan:** Data curation; formal analysis; writing – review and editing; supervision; conceptualization. **David B. Ascher:** Methodology; formal analysis; writing – review and editing; supervision; conceptualization; formal analysis.

## Funding

DBA is supported by the investigator grant from the National Health and Medical Research Council (NHMRC) of Australia [GNT1174405]. SEM, JLD and the work is supported by a NHMRC Centre for Research Excellence Pulmonary Fibrosis, a MRFF Genomics Health Futures Mission grant and a Private Philanthropic Foundation. NN is supported by an Annie Bishop PhD scholarship, JLD is also partially supported by a Select Foundation Principal Research Fellowship.

## Notes

https://biosig.lab.uq.edu.au/charactert/

### Competing Interest Statement

The authors have declared no competing interest.

