## Supplementary Figure, Supplementary Table for "CharacTERT: A Machine Learning Tool for Classifying hTERT Missense Variants"

### Supplementary Figures

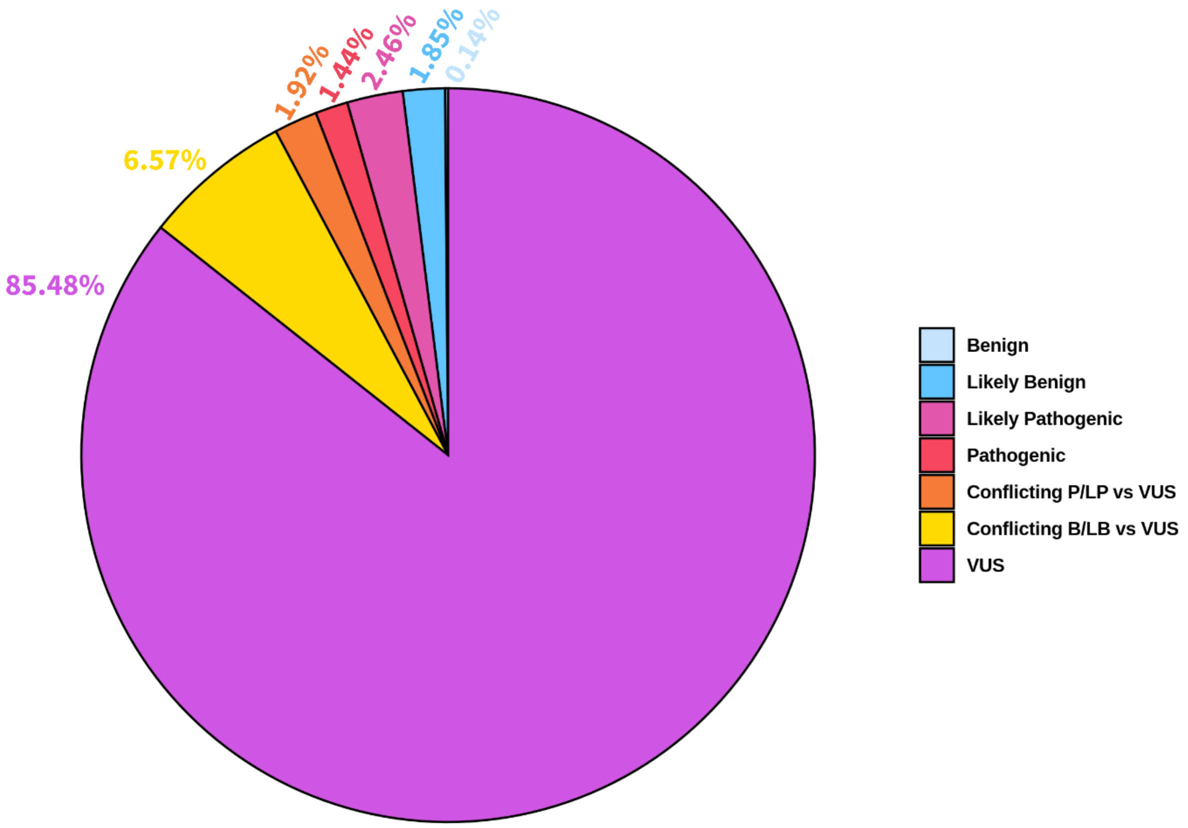


Figure S1: **Distribution of *hTERT* germline missense variants from ClinVar, accessed in February 2024.** Variants are classified as: benign, Likely benign, Likely pathogenic, pathogenic, Conflicting P/LP vs VUS, Conflicting B/LB vs VUS, and Variant of Uncertain Significance (VUS). The percentages show the proportion of each category within the total dataset, which consists of 1,464 variants.

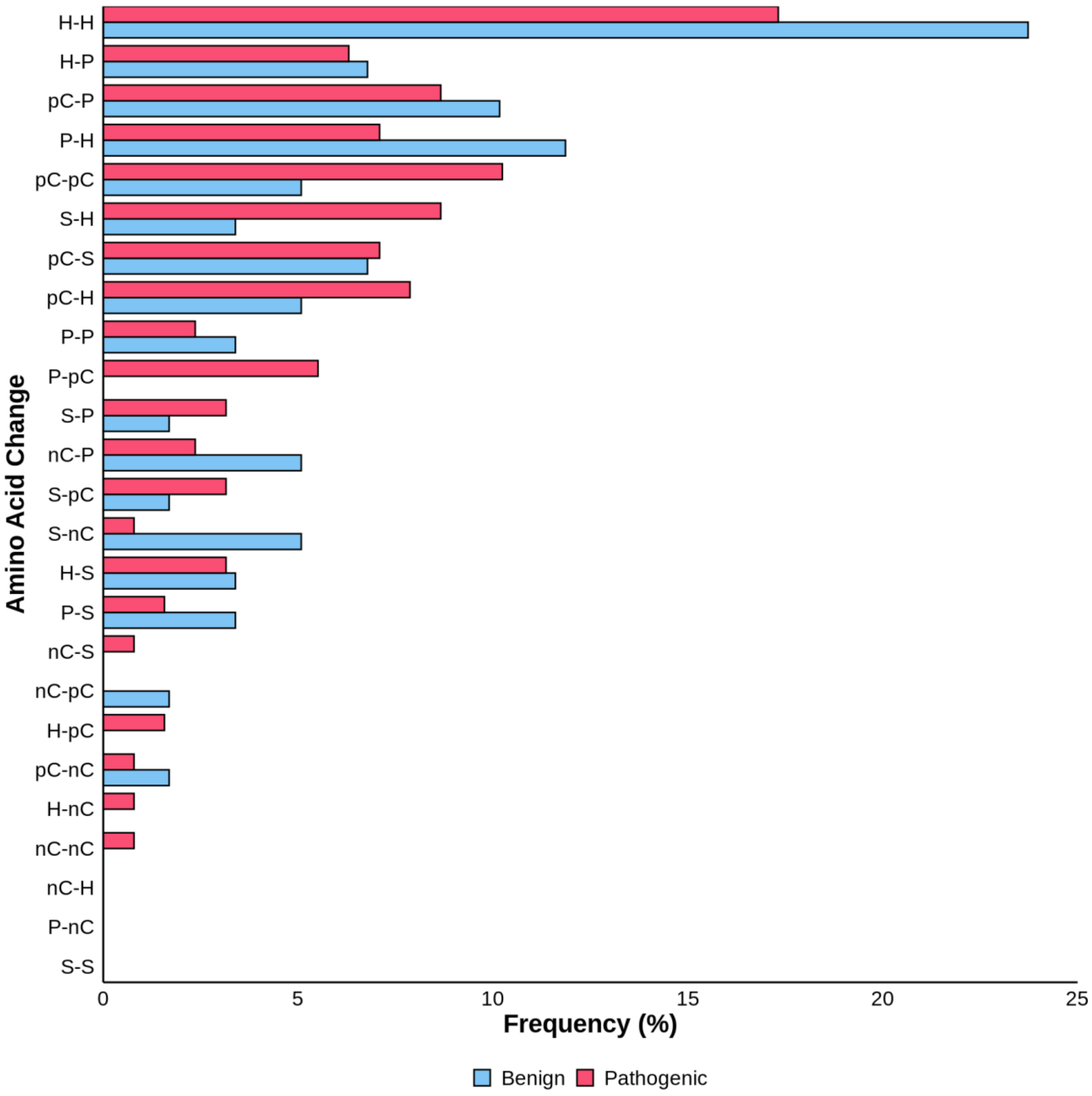


Figure S2: **Frequency of different types of amino acid substitutions in hTERT protein from the variants included in the *TERT* integrated dataset.** The 20 amino acids were categorized into five different groups considering their biochemical properties: hydrophobic (A, F, I, L, M, V, W, Y), polar (N, Q, S, T), negatively charged (D, E), positively charged (H, K, R), and special (C, G, P). These groups were then labelled as follows: hydrophobic (H), polar (P), negatively charged (NC), positively charged (PC), and special (S).

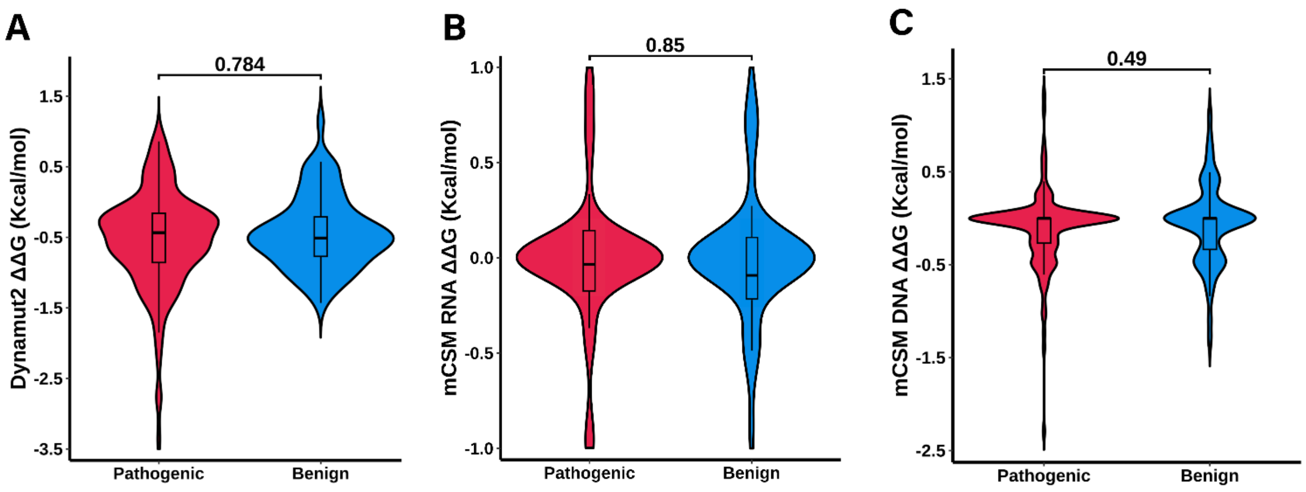


Figure S3: **Identifying potential pathogenic drivers of hTERT.** Statistical analysis was performed using the Wilcoxon rank-sum test to evaluate the mutation effect on protein stability captured by Dynamut2 (**A**), and mutation effect on both protein-RNA (**B**) and protein-single-stranded DNA (**C**) binding affinity calculated by mCSM-NA. The analysis was conducted to identify significant differences between pathogenic and benign mutations, with a significance level of 5%.

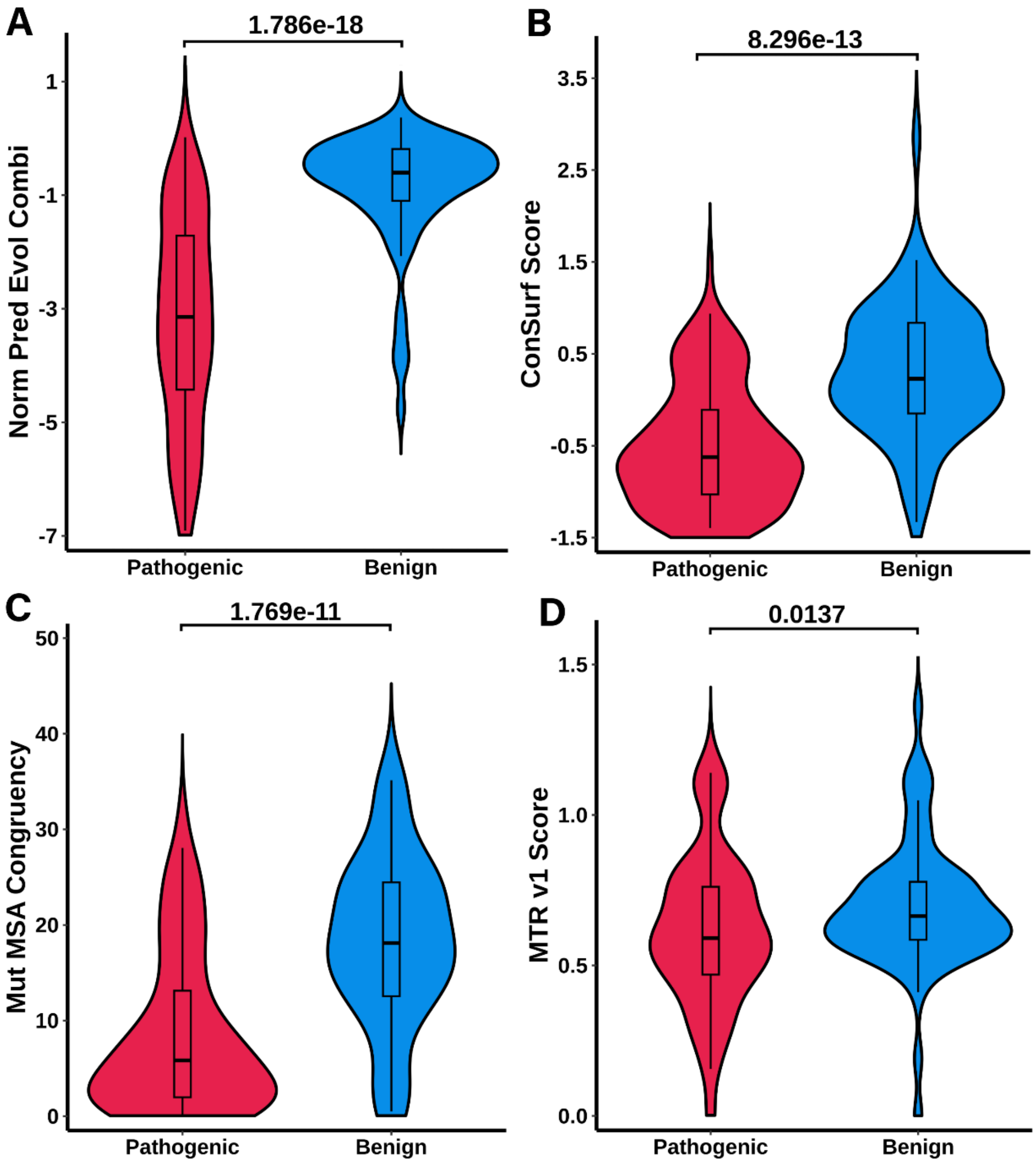


Figure S4: **Identifying potential pathogenic drivers of hTERT.** Statistical analysis was performed using the Wilcoxon rank-sum test to evaluate evolutionary conservation (**A-D**) captured by GEMME (**A**), ConSurf (**B**), and Envision (**C**), as well as population conservation (**D**) captured by MTR. The analysis was conducted to identify significant differences between pathogenic and benign mutations, with a significance level of 5%.

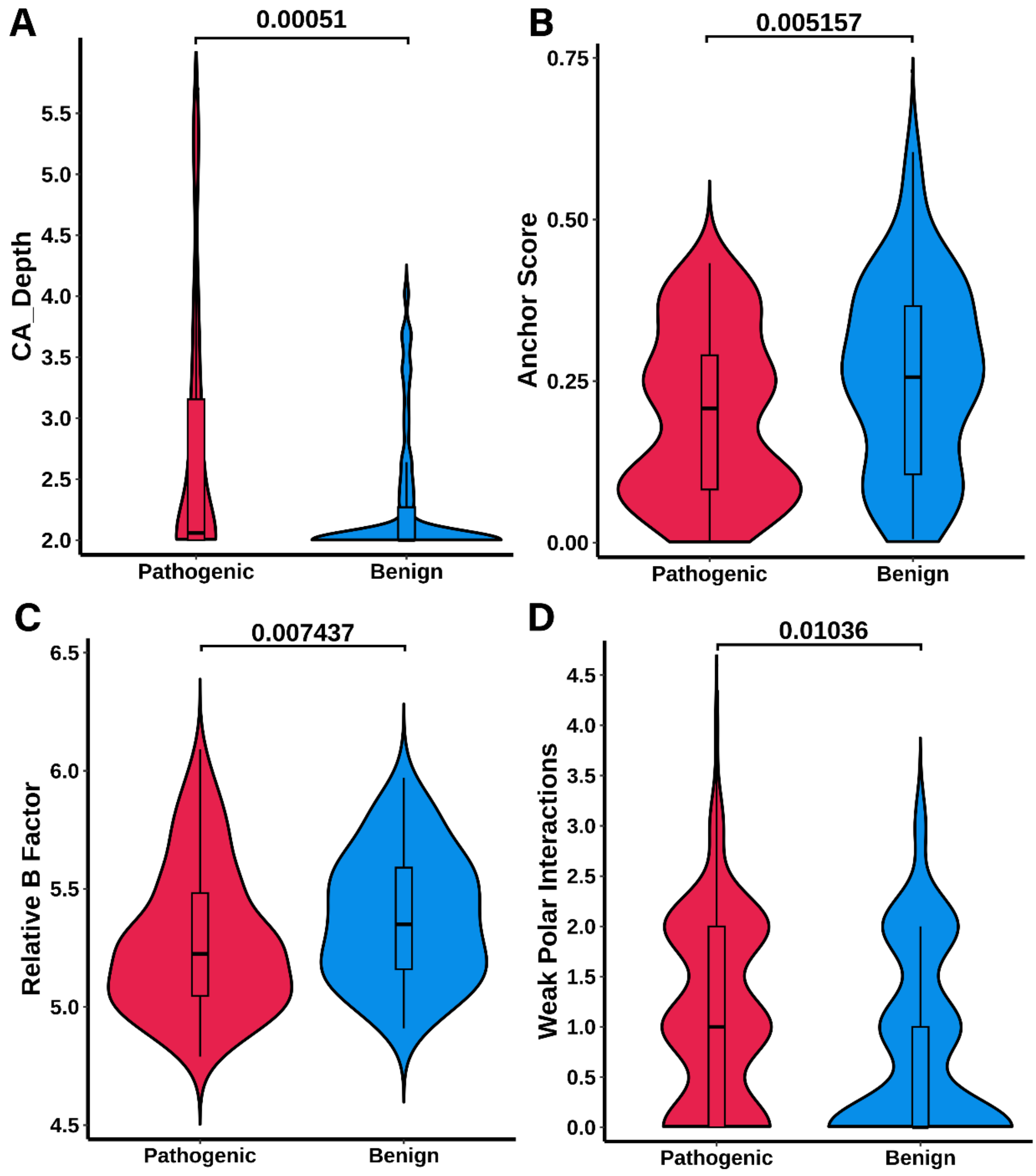


Figure S5: **Identifying potential pathogenic drivers of *hTERT*.** Statistical analysis was performed using the Wilcoxon rank-sum test to evaluate the biochemical properties of the mutation local environment (**A-D**) captured by Contact Area Depth (**A**), ANCHOR score (**B**), Relative B factor (**C**), and Arpeggio Weak Polar Interactions (**D**). The analysis was conducted to identify significant differences between pathogenic and benign mutations, with a significance level of 5%.

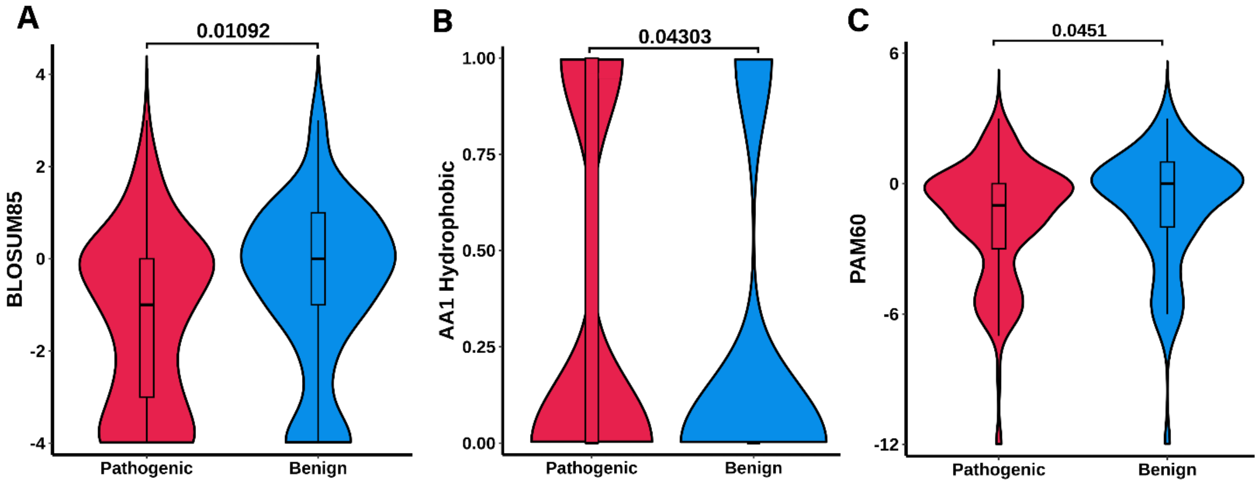


Figure S6: **Identifying potential pathogenic drivers of *hTERT*.** Statistical analysis was performed using the Wilcoxon rank-sum test to evaluate the amino acidic biochemical properties (**A-C**) captured by BLOSUM matrix (**A**), Envision (**B**), and PAM matrix (**C**). The analysis was conducted to identify significant differences between pathogenic and benign mutations, with a significance level of 5%.

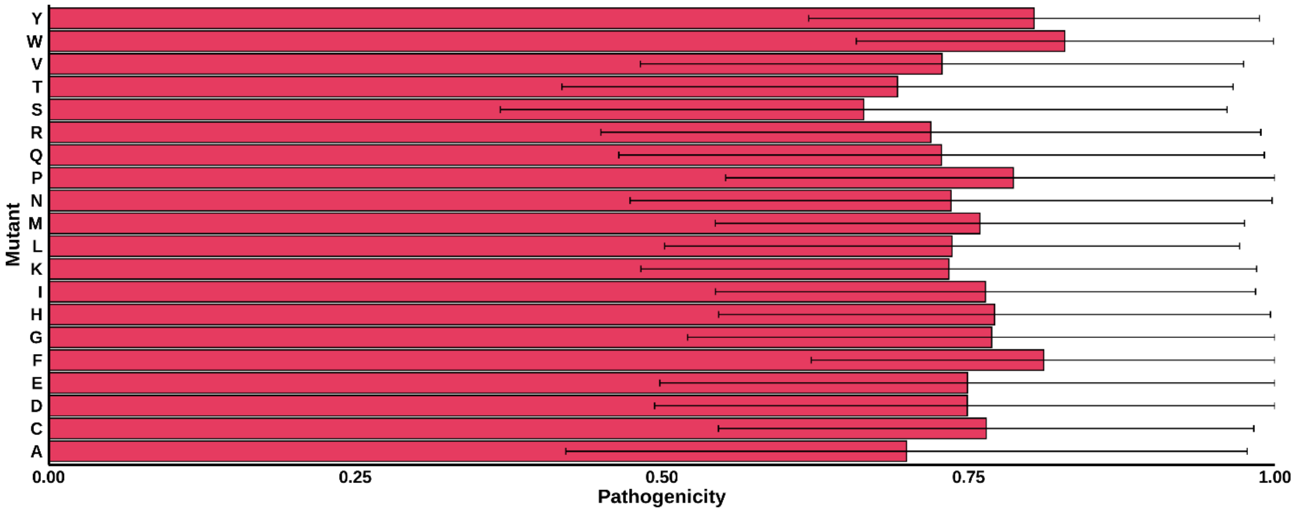


Figure S7: **Average pathogenicity of each mutant based on *in silico* saturation mutagenesis results using Position 10CV model.** Each bar represents the average pathogenicity probability predicted for the corresponding mutant, with error bars showing the standard deviation (SD) of the prediction. The x-axis displays the pathogenicity probability, ranging from 0 (benign) to 1 (pathogenic).

### Supplementary Tables

Table S1: **Qualitative analysis of features.** Features showing significant differences between pathogenic and benign mutations at a 5% significance level.

| **Feature name** | **p-value** | **Mean** | **Median** | **Standard Deviation** | **Description of feature** |
|---|---|---|---|---|---|
| Evolutionary Conservation Index (normPred_evolInd) | 4.59E-20 | -2.698 | -2.431 | 2.061 | Evolutionary conservation of the mutated position based on individual sequences, computed using GEMME (1). |
| Combined Evolutionary Index (normPred_evolCombi ) | 1.79E-18 | -2.489 | -2.148 | 1.947 | Combined evolutionary conservation of the mutated position across related sequences, computed using GEMME (1). |
| Epistasis Conservation Index (normPred_evolEpi) | 8.77E-18 | -0.423 | -0.399 | 0.296 | Conservation score reflecting epistasis effects at the mutated position, computed using GEMME (1). |
| ΔPSIC | 5.13E-15 | 1.423 | 1.244 | 1.146 | Change in Position-Specific Independent Counts score between the wild type and mutant residue, available at Envision (2). |
| Wild-Type PSIC Score | 1.96E-13 | -1.814 | -1.806 | 0.538 | PSIC score for the wild-type amino acid, available on Envision (2). |
| ConSurf Conservation Score | 8.30E-13 | -0.255 | -0.332 | 0.770 | Conservation score indicating evolutionary variability of an amino acid position, available on ConSurf (3). |
| Envision Functional Predictions | 2.26E-12 | 0.922 | 0.934 | 0.077 | Functional impact prediction scores for mutations, available on Envision (2). |
| Mutation MSA Congruency | 1.77E-11 | 11.518 | 8.937 | 9.473 | Consistency of the mutation with observed amino acid frequencies in multiple sequence |

| | | | | | |
|---|---|---|---|---|---|
| | | | | | alignments, available on Envision (2). |
| Mutant PSIC Score | 8.27E-08 | -3.237 | -2.985 | 0.955 | PSIC score for the mutant amino acid, available on Envision (2). |
| Relative Solvent Accessibility (RSA) | 6.41E-06 | 0.33 | 0.29 | 0.27 | Relative solvent accessibility of a mutation |
| MTR3D (14 Å window size) | 9.62E-06 | 0.672 | 0.668 | 0.087 | Missense Tolerance Ratio within a 14 Å structural window; available at MTR3D (4). |
| Hydrophobic | 1.04E-05 | 3.085 | 1 | 5.023 | Measure of hydrophobicity of the mutation, obtained from Arpeggio (5). |
| MTR3D (8 Å window size) | 1.64E-05 | 0.661 | 0.661 | 0.147 | Missense Tolerance Ratio within an 8 Å structural window; available at MTR3D (4). |
| Sequence Proximity to Mutation (Seq_ind_closest_mut) | 1.85E-05 | 52.423 | 52.65 | 28.251 | Sequence distance to the closest mutation in the dataset, obtained from Envision (2). |
| MTR3D (11 Å window size) | 2.66E-05 | 0.661 | 0.658 | 0.109 | Missense Tolerance Ratio within an 11 Å structural window; available at MTR3D (4). |
| mCSM- PPI ΔΔG | 0.0001633 18 | -0.463 | -0.352 | 0.564 | Predicted change in protein-protein interaction (PPI) binding free energy due to the mutation, computed using mCSM-PPI (6). |
| Proximal | 0.0004888 98 | 65.887 | 67 | 29.751 | Proximity of the mutation site to key functional regions, obtained from Arpeggio (5). |
| Cα Depth | 0.0005098 17 | 2.643 | 2 | 1.072 | Depth of the C-alpha atom of the mutated residue in the protein structure. |
| IUPRED2 Disorder Score | 0.0009983 18 | 0.142 | 0.116 | 0.120 | Intrinsic disorder prediction score for the mutated residue, |

| | | | | | |
|---|---|---|---|---|---|
| | | | | | available at IUPRED2 (7). |
| Solvent Accessibility-SST (KOSJ950100_RSA_SST) | 0.001864794 | 1.885 | 1.2 | 2.023 | Combination of relative solvent accessibility and secondary structure terms for the mutated site, obtained from Aaindex (8). |
| IUPRED3 Disorder Score | 0.002396824 | 0.140 | 0.125 | 0.114 | Intrinsic disorder prediction score for the mutated residue, obtained from IUPRED3 (7) |
| LUTR910104 | 0.002466984 | -0.756 | 2 | 12.810 | Amino acid property index for the mutated residue, obtained from Aaindex (8). |
| CarbonPI | 0.002596327 | 0.812 | 0 | 1.297 | Carbon atom properties and interactions of the mutation, computed using Arpeggio (5). |
| d_VDWClash | 0.003083361 | -1.132 | 0 | 5.489 | Van der Waals radii of the mutation computed using Arpeggio (5). |
| MTR3D (5 Å window size) | 0.003313355 | 0.660 | 0.653 | 0.236 | Missense Tolerance Ratio within a 5 Å structural window, available at MTR3D (4). |
| THOP960101 | 0.003615433 | -0.174 | -0.09 | 0.498 | Amino acid property index reflecting thermal stability, obtained from Aaindex (8). |
| QU_C930101 | 0.003735689 | 0.156 | 0.093 | 0.263 | Amino acid property index reflecting residue substitution, obtained from Aaindex (8). |
| Anchor Stability Score | 0.005157339 | 0.214 | 0.228 | 0.136 | Predicted impact on protein stability and anchor regions, computed using IUPRED3 (7) |
| Unstructured Secondary Structure (SST_None) | 0.005518788 | 0.207 | 0 | 0.406 | Secondary structure information indicating no defined structure. |
| Relative B-Factor | 0.007436682 | 5.308 | 5.27 | 0.293 | Relative B-factor indicating flexibility |

| | | | | | |
|---|---|---|---|---|---|
| | | | | | of the mutated residue. |
| ZHAC000103 | 0.0078928 16 | 0.281 | 0.47 | 0.951 | Amino acid property index reflecting hydrophobicity, obtained from Aaindex (8). |
| LUTR910109 | 0.0098937 07 | -0.432 | 2 | 10.848 | Amino acid index reflecting residue exposure, obtained from Aaindex (8). |
| MEHP950102 | 0.0101752 03 | 1.021 | 0.99 | 0.224 | Amino acid index related to hydrophobic moment, obtained from Aaindex (8). |
| BLOSUM90 | 0.0102502 39 | -1.277 | -1 | 2.02 | Substitution matrix score for amino acid changes based on evolutionary data, obtained from BLOSUM90 matrix (9,10). |
| WeakPolar | 0.0103604 46 | 0.991 | 1 | 0.927 | Less strict weak hydrogen bonding (without angle terms) computed using Arpeggio (5). |
| MTR v1 (21 codon window size) | 0.0107112 03 | 0.472 | 0.455 | 0.180 | Missense Tolerance Ratio within a 21-codon sequence window; available at MTR (11). |
| BLOSUM85 | 0.0109232 53 | -1.136 | -1 | 1.882 | Substitution matrix score for amino acid changes based on evolutionary data, obtained from BLOSUM85 matrix (9,10). |
| ZHAC000106 | 0.0127389 87 | 0.253 | 0.38 | 0.642 | Amino acid property index for hydrophobicity and interaction, obtained from Aaindex (8). |
| MTR v1 (41 codon window size) | 0.0136963 15 | 0.639 | 0.618 | 0.242 | Missense Tolerance Ratio, available on MTR (11). |
| BONM030101 | 0.0138446 51 | 0.027 | 0.1 | 0.652 | Missense Tolerance Ratio within a 41-codon sequence window; available at MTR (11). |

| | | | | | |
|---|---|---|---|---|---|
| TANS760101 | 0.019102872 | -4.781 | -4.6 | 1.158 | Amino acid index for transition frequencies, obtained from Aaindex (8). |
| BASU010101 | 0.02055436 | -0.05 | 0.002 | 0.177 | Amino acid property index for sequence analysis, obtained from Aaindex (8). |
| WEIL970102 | 0.025998902 | 0.624 | 0 | 1.321 | Amino acid property index for residue stability, obtained from Aaindex (8). |
| KOSJ950115 | 0.026451897 | 2.101 | 1.4 | 1.972 | Combination of relative solvent accessibility and flexibility indices, obtained from Aaindex (8). |
| KOSJ950100_SST | 0.02762742 | 2.218 | 1.4 | 2.075 | Combination of secondary structure and solvent accessibility indices, obtained from Aaindex (8). |
| Secondary Structure Bend | 0.028313077 | 0.099 | 0 | 0.299 | Predicted secondary structure bend for the mutation site. |
| RISJ880101 | 0.029113285 | 0.547 | 0.8 | 1.211 | Amino acid property index for isoelectric point, obtained from Aaindex (8). |
| LUTR910106 | 0.029914304 | 3.225 | 3 | 13.205 | Amino acid index for residue stability, obtained from Aaindex (8). |
| BLOSUM30 | 0.030239297 | -0.249 | 0 | 1.809 | Substitution matrix score for amino acid changes based on evolutionary data; derived from BLOSUM30 matrix (9,10). |
| BLOSUM62 | 0.030295845 | -0.624 | 0 | 1.578 | Substitution matrix score for amino acid changes based on evolutionary data; derived from BLOSUM62 matrix (9,10). |
| Observed MTR v2 (MTR_v2_obs) | 0.031153603 | 0.516 | 0.5 | 0.103 | Observed Missense Tolerance Ratio based on mutation |

| | | | | | |
|---|---|---|---|---|---|
| | | | | | data, available at MTR (11). |
| BLOSUM60 | 0.0311704 47 | -0.610 | 0 | 1.588 | Substitution matrix score for amino acid changes based on evolutionary data, obtained from BLOSUM60 matrix (9,10). |
| HENS920103 | 0.0312373 45 | -1.409 | -1 | 2.677 | Amino acid property index for hydropathy, obtained from Aaindex (8). |
| BLOSUM65 | 0.0334854 63 | -0.681 | 0 | 1.702 | Substitution matrix score for amino acid changes based on evolutionary data, obtained from BLOSUM65 matrix (9,10). |
| MUET020101 | 0.0362788 06 | -0.333 | 0 | 2.308 | Amino acid property index for residue flexibility, obtained from Aaindex (8). |
| SKOJ970101 | 0.0364058 27 | 0.010 | 0 | 0.640 | Amino acid property index for residue charge, obtained from Aaindex (8). |
| BLOSUM95 | 0.0364122 14 | -1.465 | -1 | 1.939 | Substitution matrix score for amino acid changes based on evolutionary data, obtained from BLOSUM95 matrix (9,10). |
| BLOSUM80 | 0.0377286 79 | -0.925 | -1 | 1.877 | Substitution matrix score for amino acid changes based on evolutionary data, obtained from BLOSUM80 matrix (9,10). |
| MTR v2 | 0.0413315 51 | 0.701 | 0.702 | 0.136 | Missense Tolerance Ratio based on mutation data, available at MTR (11). |
| LIWA970101 | 0.0417538 14 | -3.191 | -3.02 | 0.943 | Amino acid property index for sequence compatibility, obtained from Aaindex (8). |

| | | | | | |
|---|---|---|---|---|---|
| Wild-Type Hydrophobicity (AA1_Hydrophobic) | 0.043032585 | 0.296 | 0 | 0.458 | Hydrophobicity score for the wild-type amino acid, available at Envision (2). |
| ZHAC000102 | 0.043133991 | 0.123 | 0 | 0.416 | Amino acid property index for residue polarity, obtained from Aaindex (8). |
| KESO980101 | 0.04400385 | -3.681 | -3.75 | 1.321 | Amino acid property index for residue substitution cost, obtained from Aaindex (8). |
| PAM60 | 0.045099125 | -1.460 | -1 | 2.726 | Substitution matrix score for amino acid changes based on evolutionary data, obtained from PAM60 matrix (12). |
| Mutant Polarity (AA2_Polar) | 0.045612657 | 0.249 | 0 | 0.433 | Prediction of polarity impact for the mutant amino acid, available on Envision (2). |

Table S2: **Leave-One-Group-Out cross validation performance.** Domain 4CV predictive performance in the training set, measured by Matthew's Correlation Coefficient (MCC), accuracy, sensitivity, and specificity.

| Domain left Out for Testing | MCC | Accuracy | Sensitivity | Specificity |
|---|---|---|---|---|
| NTE | 0 | 0.86 | 0.86 | 0 |
| RBD | 0.47 | 0.75 | 0.82 | 0.64 |
| RT | 0.73 | 0.88 | 0.93 | 0.79 |
| CTE | 0.86 | 0.94 | 0.96 | 0.9 |

Table S3: **Misclassified variants.** List of the misclassified variants by the three different models, in the training and blind test sets, along with their corresponding ΔΔE scores.

| Variants | Set | Predicted Label | Actual Label | ΔΔE scores |
|---|---|---|---|---|
| A67V | Training | Benign | Pathogenic | -0.37 |
| R342Q | Blind Test | Pathogenic | Benign | -3.46 |
| G527V | Training | Pathogenic | Benign | -2.07 |
| D628N | Training | Benign | Pathogenic | -0.52 |
| A670V | Training | Pathogenic | Benign | -1.13 |
| P703L | Training | Pathogenic | Benign | -1.43 |
| P785L | Training | Benign | Pathogenic | -0.98 |
| N906S | Training | Pathogenic | Benign | -4.76 |

Table S4: **Misclassified variants by Position 10CV.** List of the misclassified variants by the Position 10CV model, in the training and blind test sets.

| Variants | Set | Predicted Label | Actual Label | ΔΔE scores |
|---|---|---|---|---|
| P33S | Training | Benign | Pathogenic | -1.03 |
| A67V | Training | Benign | Pathogenic | -0.42 |
| P322L | Training | Pathogenic | Benign | -0.89 |
| R342Q | Blind Test | Pathogenic | Benign | -3.04 |
| P380S | Training | Pathogenic | Benign | -3.13 |
| R522K | Training | Benign | Pathogenic | -0.05 |
| V526A | Training | Pathogenic | Benign | -0.05 |
| G527V | Training | Pathogenic | Benign | -1.81 |
| T567M | Training | Benign | Pathogenic | -0.80 |
| R595K | Training | Pathogenic | Benign | -0.68 |
| Q608H | Training | Pathogenic | Benign | -1.01 |
| D628N | Training | Benign | Pathogenic | -0.46 |
| T644M | Training | Pathogenic | Benign | -2.01 |
| E652K | Training | Pathogenic | Benign | -1.35 |
| V664L | Training | Benign | Pathogenic | -0.55 |
| A670V | Training | Pathogenic | Benign | -1.10 |
| R671Q | Training | Pathogenic | Benign | -0.73 |
| R672H | Training | Pathogenic | Benign | -0.98 |
| P703L | Training | Pathogenic | Benign | -1.41 |
| E705Q | Training | Benign | Pathogenic | -0.32 |
| P785L | Training | Benign | Pathogenic | -0.88 |
| A817T | Training | Pathogenic | Benign | -1.38 |
| N906S | Training | Pathogenic | Benign | -4.72 |
| R962C | Training | Pathogenic | Benign | -0.83 |
| R962H | Training | Pathogenic | Benign | -0.71 |
| S1067P | Training | Benign | Pathogenic | -0.51 |
| R1086C | Training | Benign | Pathogenic | -0.88 |
| T1110M | Training | Benign | Pathogenic | -0.45 |
| E1116Q | Training | Benign | Pathogenic | -0.25 |

Table S5: **Residues in the linker region predicted as Pathogenic.** List of the residue positions which the Position 10CV model predicted to be pathogenic.

| Position | Average Confidence |
|---|---|
| W203 | 0.63 |
| R208 | 0.51 |
| L216 | 0.50 |
| G220 | 0.53 |
| R222 | 0.86 |
| R223 | 0.93 |
| R224 | 0.69 |
| S231 | 0.51 |
| P233 | 0.62 |
| K236 | 0.74 |
| R237 | 0.71 |
| P238 | 0.63 |
| R239 | 0.81 |
| E245 | 0.61 |
| P246 | 0.54 |
| G252 | 0.51 |
| S255 | 0.65 |
| A257 | 0.59 |
| F270 | 0.65 |
| V273 | 0.64 |
| S274 | 0.56 |

| Position | Average Confidence |
|---|---|
| P275 | 0.52 |
| A276 | 0.61 |
| R277 | 0.51 |
| C279 | 0.54 |
| E280 | 0.54 |
| S284 | 0.62 |
| E286 | 0.52 |
| G287 | 0.57 |
| L289 | 0.56 |
| P297 | 0.60 |
| R301 | 0.55 |
| Q302 | 0.57 |
| P308 | 0.50 |
| S309 | 0.56 |
| R312 | 0.66 |
| P314 | 0.58 |
| P316 | 0.77 |
| W317 | 0.66 |
| D318 | 0.57 |
| T319 | 0.54 |
| P320 | 0.64 |
